# Investigating the right heart hemodynamics in the Tetralogy of Fallot: a computational study

**DOI:** 10.1101/2024.09.13.612742

**Authors:** Francesca Renzi, Giovanni Puppini, Giovanni B. Luciani, Christian Vergara

## Abstract

Characterizing flow within the right heart (RH) is particularly challenging due to its complex geometries. However, gaining insight into RH fluid dynamics is of extreme diagnostic importance given the high prevalence of acquired and congenital heart diseases with impaired RH functionality. In this study, we present a comprehensive, patient-specific, image-based computational analysis of the hemodynamics of the RH in healthy and repaired-ToF (ToF) cases. From multi-series cine-MTRI, we reconstruct the geometry and motion of the patient’s right atrium, ventricle, and pulmonary and tricuspid valves. For this purpose, we develop a novel and flexible reconstruction procedure that enables us, for the first time, to integrate fully patient-specific tricuspid valve dynamics into a computational model, enhancing the accuracy of our RH blood flow simulations. This work provides novel insight into the altered hemodynamics of repaired-ToF RH with severe pulmonary regurgitation and into the hemodynamics changes induced by the pulmonary valve replacement intervention. Modelling the whole RH enables us to understand the contribution of the superior and inferior vena cava inflows to the ventricular filling, as well as the impact of the impaired right atrial function on the ventricular diastole. We analyze the turbulent and transitional behaviour by including the Large-Eddies Simulation sigma model in our computational framework. This highlights how the contribution of the smallest scales in the dissipation of the turbulent energy changes among health and disease.

## 1 Introduction

The right heart (RH) significantly impacts many congenital and acquired heart disorders (Dumitrescu et al., 2018). Among all, the Tetralogy of Fallot (ToF) is the most common cyanotic congenital heart disease, occurring in approximately 1 in 3’500 births and accounting for about 10% of all congenital malformations (Valsangiacomo Buechel and Mertens, 2012). Infants affected by such pathology undergo a reparative surgery in their first months of birth in order to restore the correct blood oxygenation (*repaired-ToF*). Although the survival rate after the intervention has remarkably improved over the years (Cuypers et al., 2014), repaired-ToF patients may develop several complications in the adult age, such as *pulmonary regurgitation* (PR), bi-ventricular dysfunction, and arrhythmias (Redington, 2006). These dramatically hamper the life quality, leading to RH dysfunction and eventually heart failure, calling for the need for further surgical interventions in the adult age, e.g. through *pulmonary valve replacement* (PVR). In particular, among all the late outcomes, PR stands out as the most important, as it has been shown to be linked to right ventricle (RV) remodelling processes (Redington, 2006; Senthilnathan et al., 2013). For these reasons, the appropriately timed PVR intervention is of utmost importance in halting the RV remodelling and allowing RH function recovery. Nowadays, the main tools for determining RV remodelling and PR are cardiac magnetic resonance imaging (cMRI) and echocardiography. However, both technologies do not provide a comprehensive analysis of intracardiac flow pre- and post-PVR (Senthilnathan et al., 2013). In contrast, computational fluid dynamics (CFD) models allow for the detailed measurement of hemodynamics alteration following such intervention, offering a valuable understanding of the surgical outcomes in both the short and long term.

Developing in-silico models for the RH of repaired-ToF patients is a particularly challenging task due to the need to account both for the variability of the cardiac morphology and for the specific cardiac wall movement. The first requirement is met by building patient-specific computational models, which provide insight into local cardiac hemodynamics and have proven informative in addressing various queries in other clinical contexts as well (Biglino et al., 2017; Fumagalli et al., 2024). To address the second requirement, the literature presents two approaches that differ in how they recover the cardiac chamber’s displacement. In the case of *fluid-structure interaction* (FSI) models, the displacement is an unknown of the problem (Bucelli et al., 2023; Viola et al., 2022). These models are generally characterized by high computational costs, require meticulous calibration of the myocardial properties, and need to recover the unloaded configuration. In the second approach, the displacement can be prescribed either from offline electromechanical simulations (Augustin et al., 2016; Zingaro et al., 2024) or from images. In the latter case, we talk about *dynamic image-based CFD* (DIB-CFD) models (Bennati et al., 2023b; Chnafa et al., 2016; Fumagalli et al., 2022; Seo et al., 2014; Wiputra et al., 2016; Mangual et al., 2012; Collia et al., 2021). Despite the main drawback of complex image processing needed to recover a meaningful and artefact-free cardiac movement, this approach has become a valid alternative to FSI model when detailed dynamical images are available. Regarding the study of ToF, both FSI (Tang et al., 2010) and DIB-CFD models (Wiputra et al., 2018; Loke et al., 2022, 2024) have been recently developed for the study, e.g., of turbulence and stresses characterizing such condition.

All the DIB-CFD works regarding the RH focused so far on the ventricular fluid dynamics solely. However, for the accurate and comprehensive analysis of the diastolic phase, the inclusion of the atrium is fundamental, e.g., to provide the right initialization for the systolic phase. Furthermore, these studies neglected the presence of (and thus the interaction with) pulmonary and tricuspid valves (PV and TV). This omission could represent a significant limitation as blood dynamics is highly influenced by the interaction with valves, especially in pathological cases and in the context of PVR. In the direction of overcoming these limitations, a first step was performed in Renzi et al. (2023), where we presented a novel technique for RH reconstruction from multi-oriented cine-MRI. This technique was then tested in a complete RH, where, however, only the systolic phase was considered, and the valve dynamics was approximated by an on/off behaviour.

In the present study, we want to overcome the previous limitations by developing a comprehensive DIB-CFD model based on time-resolved cine-MRI of the whole right heart (RV+RA+TV+PV). Specifically, we simulate the whole heartbeat, including the patient-specific dynamics of the valves and their interaction with blood. Thanks to these new elements, we provide a deep analysis of the hemodynamic changes that occur immediately after the implantation of a competent PV in severely regurgitant repaired-ToF patients. For this purpose, we study the hemodynamics in three different scenarios, namely, a healthy and two repaired-ToF cases.

In this respect, the main contributions of this work are:

1. The introduction of a novel methodology for the reconstruction of tricuspid valve configuration and dynamics from arbitrarily oriented cine-MRI;
2. The development of fully RH patient-specific DIB-CFD simulation for the whole heartbeat with imposed motion of ventricle, atrium, pulmonary artery root, pulmonary and tricuspid valves, all reconstructed from cine-MRI;
3. The investigation of hemodynamics changes right after PVR implantation;
4. The analysis of blood-stagnation related quantities (e.g., blood residence time) which were never investigated in the context of RH;
5. The introduction of new insights into the comparison between the right and left hemodynamics.

The new aspects of this research enable us to bring forth an original computational analysis for the repaired-ToF and to underscore the suitability of the reconstruction method proposed in Renzi et al. (2023) for DIB-CFD modelling.

## 2 Methods

In this section, we detail the image-processing procedure for the reconstruction of the patient-specific RH geometries and tricuspid and pulmonary valves, and we present the mathematical model for the description of the blood flow within the whole RH and its interaction with the endocardial walls and the valves. Finally, we introduce the quantities of interest obtained by post-processing the numerical results.

### 2.1 Available clinical data

Clinically indicated and ad hoc cardiac cine-MRI studies were performed for two subjects at the Division of Radiology, University Hospital Verona, Verona, Italy. In particular, we have at disposal images for:

- a healthy subject, hereafter referred to as H;
- a patient with chronic pulmonary valve insufficiency (regurgitant fraction, RF, equal to 38%) after the repair of tetralogy of Fallot in infancy (hereafter referred to as ToF).

Ethical review board approval and informed consent were obtained from both subjects. The acquisitions were performed using Achieva 1.5T (TX) -DS (Philips, Amsterdam, Netherlands) technology and consist of slices with homogeneous in-plane spatial resolution ranging from 1.15 to 1.25 *mm* and thickness ranging from 5 to 8 *mm*. All series have a time resolution of 30 frames/cardiac cycle. In particular, the dataset is composed of the following cine-MRI series: short-axis (SA), 4 chamber long-axis (4Ch-LA), 3 chambers-LA (3Ch-LA), and 2 chambers-LA (2Ch-LA). In addition, for H, we also have rotational images over the tricuspid valve (TV-R) at our disposal.

This dataset enables the complete reconstruction of the right atrium (RA), right ventricle (RV), right ventricle outflow tract (RVOT), main pulmonary artery (PA), as well as the tricuspid (TV) and pulmonary (PV) valves. We refer the reader to Renzi et al. (2023) for a detailed description of the used dataset.

### 2.2 Reconstruction of right heart geometry and motion

In this study, we obtain the geometries of RH internal walls and their motion over the cardiac cycle employing the *MSMorph* procedure presented in Renzi et al. (2023) and specifically developed for the generation of patient-specific RH configurations from differently oriented cine-MRI acquisitions. Fig. 1 shows a schematic representation of the whole pipeline. Briefly, we segment the right atrial and ventricular endocardia on each slice of all cine-MRI series at disposal for every acquired time instant. The segmentation is performed in a semiautomatic fashion by applying a non-rigid B-Spline-based image registration algorithm implemented in the open-source library SimpleElastix (https://simpleelastix.github.io/). Then, we collect the segmented contours referring to the same time instant to obtain *N* sets of atrial and ventricular endocardium contours, being *N* the total number of frames of the cine-MRI acquisitions (see Fig. 1 -Step 1). In the second step, we deform a template surface over each set of contours to recover two separate representations of the atrial and ventricular endocardial surfaces (Fig. 1 -Step 2). Then, we compute the RA and RV deformation fields obtained with respect to the reference end-systolic configuration by employing another non-rigid B-Spline-based registration algorithm in SimpleElastix (Fig. 1 -Step 3).

**Fig 1.**
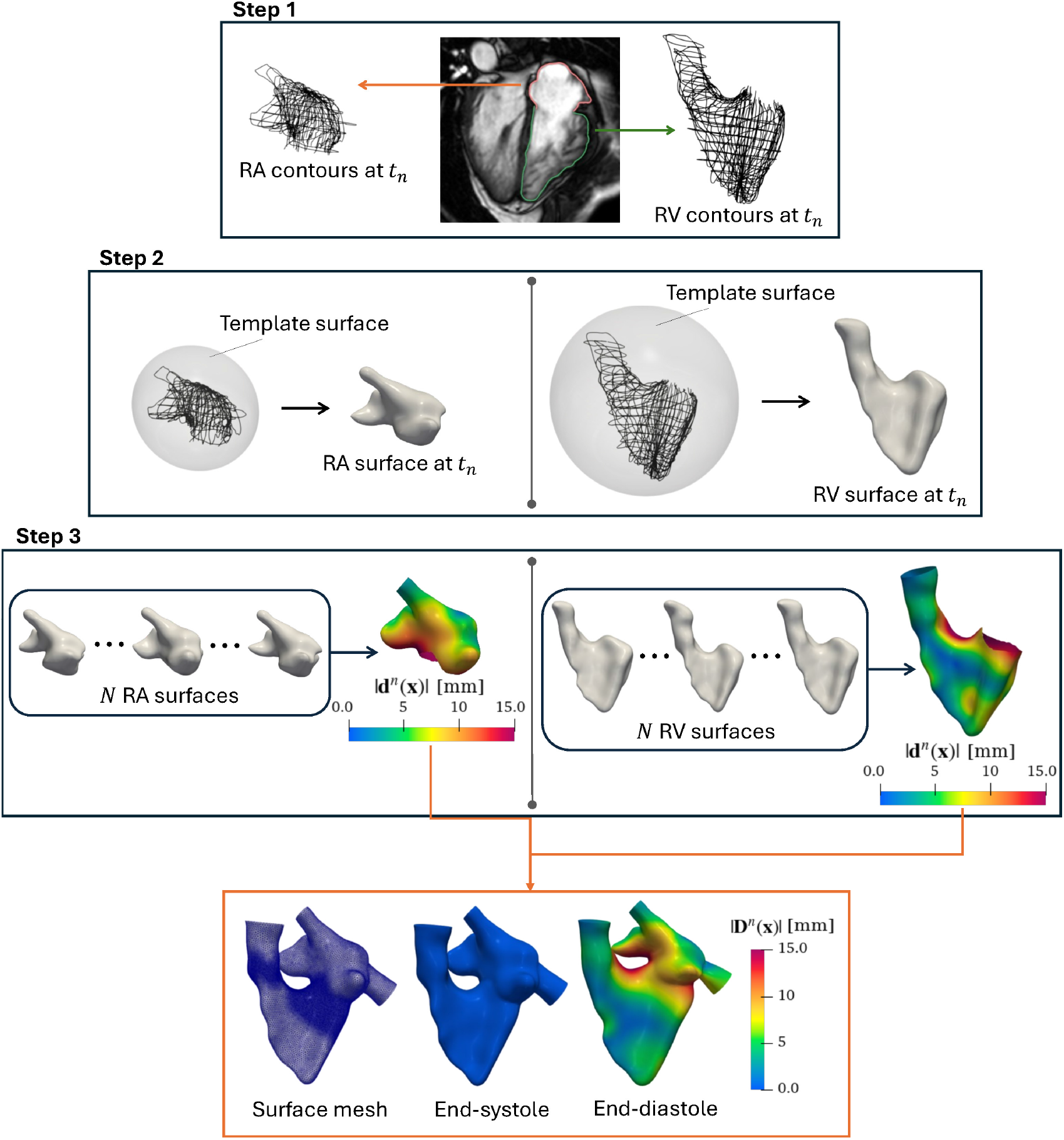
Flowchart of the reconstruction procedure for the RH (Section. 2.2). Step 1: representation of the endocardial segmentation at frame *t*_*n*_; Step 2: representation, at frame *t*_*n*_, of the RA and RV endocardial surface generated starting from a template and the segmented contours; Step 3: representation of the generation of the RA and RV displacement, **d**^*n*^(**x**), with respect to a reference configuration, starting from the *N* surfaces (output of Step 2); the magnitude of the computed displacement at the end-diastolic frame is showed; Orange box: final results of the procedure: RH surface mesh obtained for the reference end-systolic configuration (left), application of the displacement field, **D**^*n*^, computed for the end-systolic (centre) and end-diastolic (right) frame.

Finally, we built the surface mesh at the reference end-systolic configuration, starting from the connection of RA and RV surfaces. This allows us to obtain the meshes at each frame *n* by applying the deformation fields previously computed and suitably extended to the TV annulus (see Fig. 1 bottom panel).

All these geometric preprocessing steps were obtained with the software Vascular Modeling Toolkit (vmtk, http://www.vmtk.org/, Antiga et al. (2008)), enriched by additional tools for the cardiac surface processing (Fedele and Quarteroni, 2021).

### 2.3 Reconstruction of tricuspid valve geometry and motion

In this section, we present a new procedure for the generation of patient-specific TV geometry and motion, outlined in Fig. 2. The main novelty of our procedure is that it does not rely on rotational acquisitions, such as rotational cine-MRI (Stevanella et al., 2011; Bennati et al., 2023b) or transthoracic real-time three-dimensional echocardiography (Votta et al., 2008), it rather integrates informations from whatever cine-MRI series one has at its disposal where the valve is visible, by exploiting the deformation of a template.

**Fig 2.**
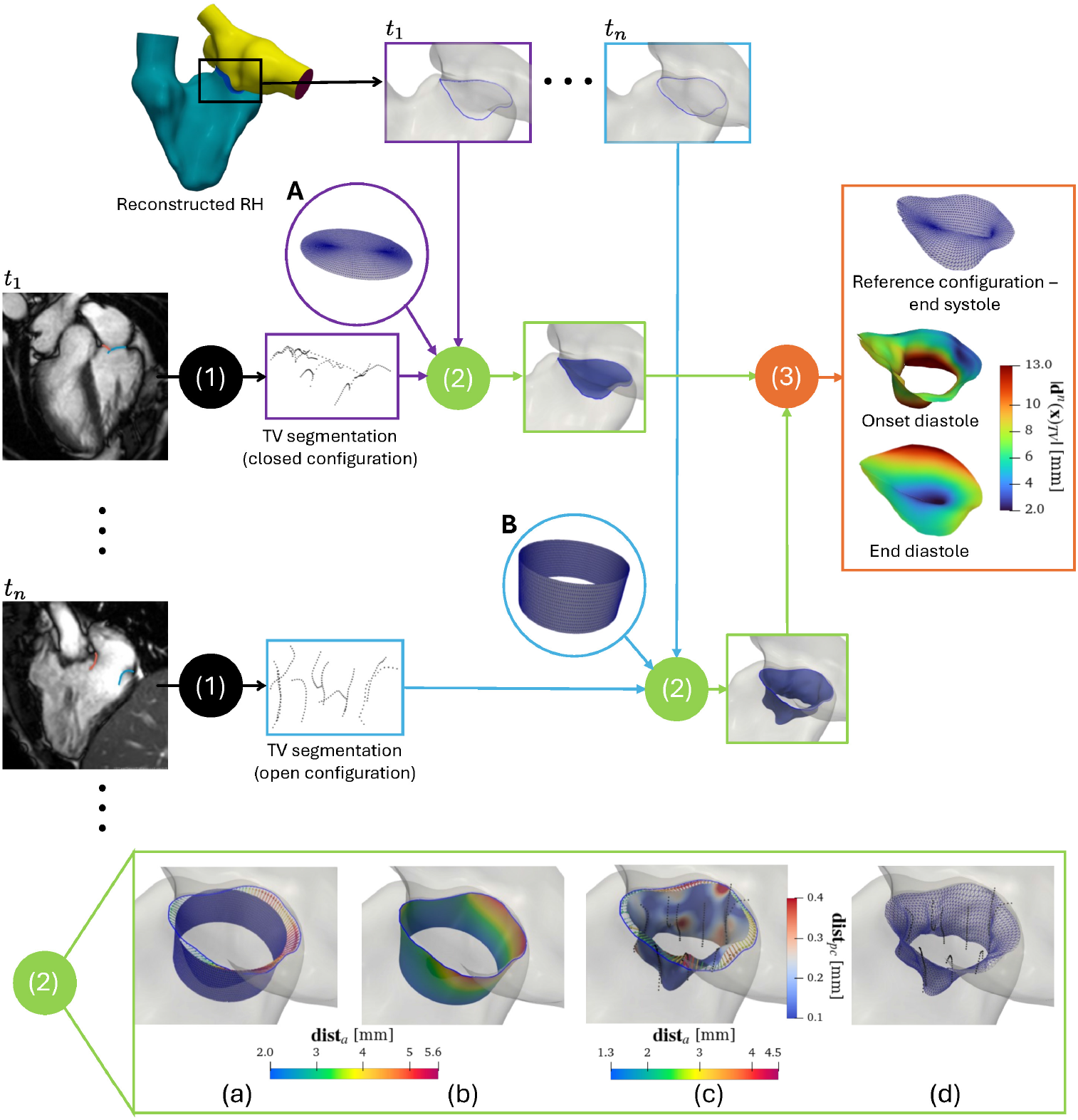
Flowchart of the reconstruction procedure for the TV (Section. 2.3) depicted at two representative instants: the onset diastole (light blue boxes and arrows) and end-diastole (purple boxes and arrows). Top row: extraction of the annulus from the RH geometry for all frames. Full circle (1): leaflets segmentation from cine-MRI data for all frames. Circle panels A and B: representation of the template surfaces for recovering the closed and open TV configurations, respectively. Full circle (2): template morphing procedure. Full circle (3): TV displacement procedure. Orange box: representation of the TV mesh at the reference end-systolic configuration (top) and representation of the displacement field magnitude at the onset (middle) and at the end (bottom) of diastole. Bottom row -green panel, focus on one iteration of the template morphing procedure: a) computation of the distance between the annulus border on RH and the annulus ring on the template, **dist**_a_; b) representation of the extended **dist**_*a*_ on the template surface deformed accordingly; c) representation of the extended distance field computed between the template surface output of b) and the segmented point cloud (black dots), **dist**_pc_, on the template surface deformed accordingly. The newly computed **dist**_*a*_ field is represented too; d) final result of the current iteration.

In particular, to recover the patient-specific TV leaflets’ morphology and motion over the heartbeat, the proposed procedure comprises the following steps which are depicted in Fig. 2:

1. The segmentation of the TV leaflets on the available cine-MRI;
2. The deformation of a template surface over the segmented TV;
3. The computation of the TV displacement field over the heartbeat.

In step 1 (see Fig. 2), for each frame, we manually trace the TV leaflets’ profile on the slices of the subset of the cine-MRI series where the valve is visible. This is accomplished by exploiting the *Markups* module of the open-source software 3D Slicer (https://www.slicer.org/). Then, we collect contours traced at the same frame together to obtain *N* separate point clouds, representing the segmentation of the subject’s valve at each frame. We notice that these segmentations are highly sparse since the valve is usually visible in a small subset of the available cine-MRI dataset.

Step 2 (depicted in the green panel of Fig. 2 for a diastolic frame) consists of i) the inclusion of a template surface into RH endocardial reconstruction and ii) the deformation over the TV segmentations, obtained in step 1, to retrieve the leaflet’s morphology. In particular, to attain i), we first compute the distance field between the TV annulus of the RH endocardium reconstruction and the closest border of the template. Then, we extend this distance field over the entire template surface and deform it accordingly. This allows us to connect the template surface to the TV annulus. After that, in ii), we compute the pointwise distance between the connected surface and the TV point cloud, we extend this field over the entire surface and warp it accordingly. In order to let the surface better resemble the segmented leaflet profiles, we iterate ii) a few times (e.g., two or three). Finally, we repeat i) to compensate for a possible surface detachment from the TV annulus and recover the connection. We observe that, as a template for step 2, we employ two different surfaces sharing the same connectivity: a disk to recover the TV closed configurations and a hole cylinder to recover the TV open configurations (see the purple circles in Fig. 2). Consequently, the connectivity is preserved among all the *N* TV surfaces, representing the valve’s time evolution.

This feature is exploited in step 3 to trivially recover the TV displacement field. Namely, we compute the displacement of each configuration with respect to a reference one (e.g. at the end-systolic instant) by simply subtracting the node coordinates of their triangulations.

The outcome of the presented procedure is the triangulated TV surface at the reference frame together with the definition of its displacement fields **d**^*n*^(**x**), *n* = 1, 舰, *N*. In Fig. 2, orange panel, we reported the reference surface and magnitude of **d**^*n*^(**x**) at two representative frames.

### 2.4 Reconstruction of pulmonary valve geometry and motion

Concerning the Pulmonary valve (PV), the cine-MR images at our disposal do not allow for a complete reconstruction of the patient-specific leaflets for any frame of the cardiac cycle. For this reason, we employ the procedure followed by Bennati et al. (2023a). Briefly, as a first step, we obtained the fully closed and fully open PV configurations by adapting the PV geometry from the *Zygote solid 3D heart model* (https://www.zygote.com) to the reconstructed patient’s PV annulus.

Starting from this configuration, we build the healthy and the following two repaired-ToF scenarios:

i. A repaired-ToF with severe pulmonary regurgitation (PVReg-ToF);
ii. A post-operative non-regurgitant scenario, obtained from case i), by considering a virtual transcatheter PVR (PVnoReg-ToF).

In order to replicate the PV insufficiency for the PVReg-ToF scenario, we further modify PV template’s closed configuration by partially closing the leaflets to induce the same regurgitant franction (RF) as the patient.

In the next step, we recover the opening and closure PV dynamics by interpolating the adapted fully open and fully (or partially) closed configurations in time. This interpolation is, in practice, realized within the Finite Elements solver (see Sect. 2.5.1).

### 2.5 Dynamic Image-Based computational fluid dynamics model

#### 2.5.1 Modeling details

Starting from the RH surfaces reconstructed as described in Sect. 2.2 at the end-systolic instant, we complete the computational domain for the DIB-CFD models of the three scenarios introduced in Sect. 1 by including the reconstructed valves (see Sect. 2.3 and 2.4). We then generate a tetrahedral volumetric mesh.

To summarize, in each scenario, the reference RH mesh, together with its valves, defines our computational domain in the reference configuration 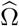. In Fig. 3 -a, we illustrate DIB-CFD domain. Here, Γ_TV_ and Γ_PV_ are the surfaces representing the TV and PV’s leaflets, respectively. By Σ_w_, we denote the boundary comprising the ventricular and atrial endocardia and the pulmonary artery, IVC, and SVC walls. The outlet section of the pulmonary artery is indicated as Σ_out_, and the inlet sections of IVC and SVC as Σ_in_.

**Fig 3.**
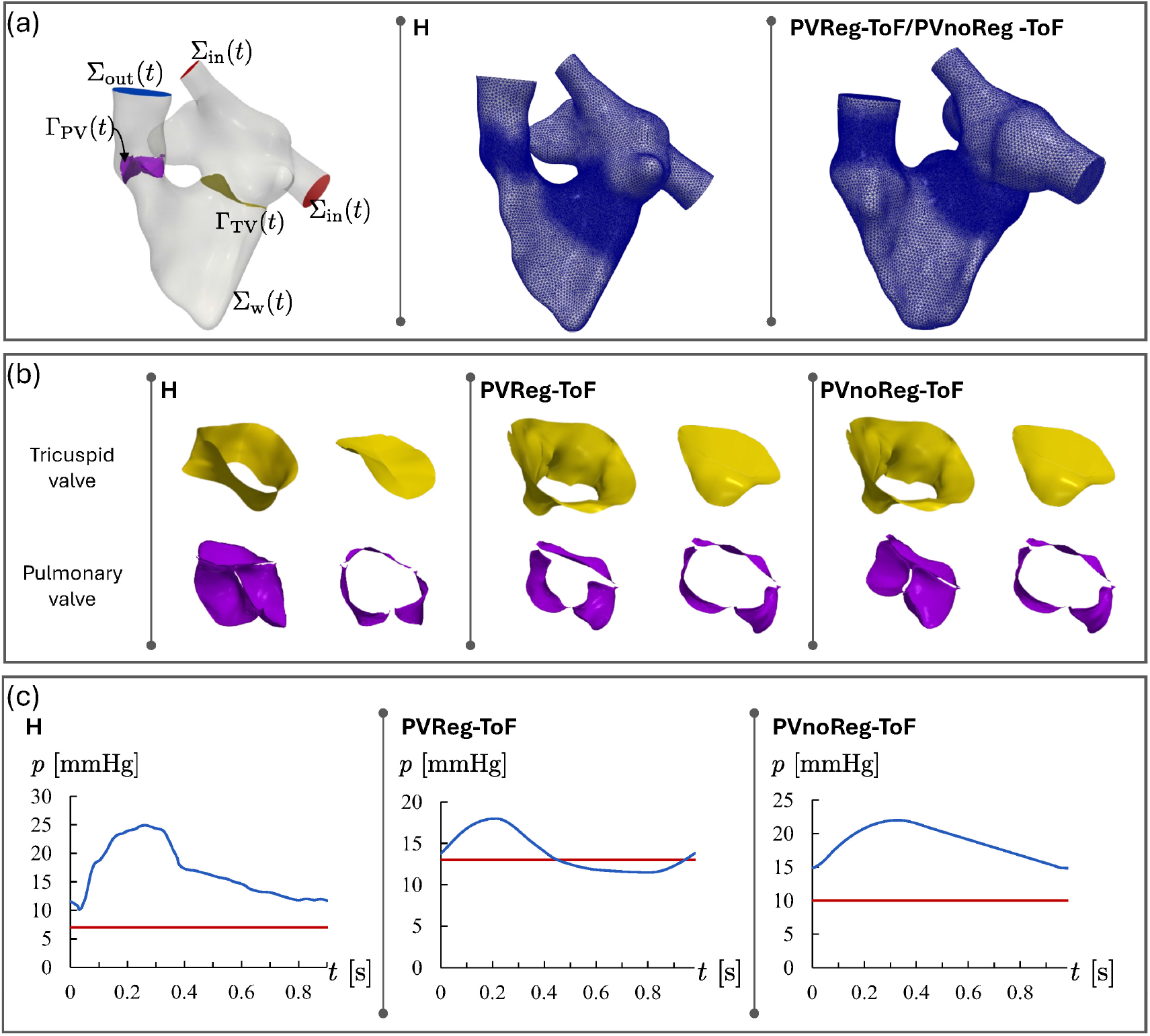
Computational domains and boundary conditions. a): on the left column the representation of the whole right heart Ω(*t*) with its boundaries showed only for the healthy (H) scenario: ventricle outflow Σ_out_(*t*) (blue); atrial inflows at the superior and inferior vena cavae Σ_in_(*t*) (red); ventricular and atrial endocardium and physical wall of the pulmonary artery Σ_w_(*t*); surface of the pulmonary valve’s leaflets Γ_PV_(*t*); surface of the tricuspid valve’s leaflets Γ_TV_(*t*). On the central and right columns, RH tetrahedric volumetric mesh for the healthy subject (H) and the repaired-ToF patient (ToF), respectively. b): tricuspid and pulmonary valves in fully open and fully closed configurations for the healthy (H), the regurgitant repaired-ToF (PVReg-ToF), and the non-regurgitant repaired-ToF (PVnoReg-ToF) scenarios. c): pressure curves at the pulmonary artery (blue line) and inferior and superior vena cave (red line), imposed as boundary conditions; values for H come from literature (Grignola, 2011; Pappano and Wier, 2018); values for PVReg-ToF and PVnoReg-ToF come from calibrated lumped parameter model (Criseo et al., 2024).

In what follows, we detail the mathematical model describing the blood flow and its interaction with the endocardium and valves in a moving setting.

We assume that blood behaved as an incompressible Newtonian fluid in heart chambers and large vessels (Quarteroni et al., 2019) with a density of 1.06 10^3^ · *kg/m*^3^ and dynamic viscosity of 3.5 · 10^−3^ *Pa* · *s*. Therefore, we consider the Navier-Stokes equations in a moving domain. As described in Sect. 2.2, the displacement **D**^*n*^(**x**) of the domain Ω(*t*) with respect to its reference configuration 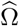was reconstructed from cine-MRI only for the acquired *N* frames. Hence, we interpolate it in time with splines to cover the whole heartbeat duration. This leads to the definition of **D**(**x**, *t*) for all *t* ∈ [0, *T*] where *T* indicates the subject’s heartbeat duration. **D**(**x**, *t*) is then used to compute the wall velocity to prescribe to the fluid problem that is solved in an Arbitrary Lagrangian-Eulerian (ALE) formulation (Hirt et al., 1997; Nobile and Formaggia, 1999). According to this framework, at each *t*, **D**(**x**, *t*) (which ia defined on Σ_w_) is extended within Ω\l “ by solving a Linear Elasticity problem (Stein et al., 2003). See Africa et al. (2024) and Renzi et al. (2023) for further details on the used solver. To account for possible transition to turbulence, we consider the Large Eddy Simulation *σ* model (Nicoud et al., 2011). Regarding the valves, we model them as thick surfaces immersed in the fluid dynamics domain without relying on fluid-structure interaction modelling. In particular, their presence is managed by the Resistive Immersed Implicit Surface (RIIS) method (Fernández et al., 2008; Fedele et al., 2017; This et al., 2020). We handle differently the valve dynamics in the cases of TV and PV. For the TV motion, we linearly interpolate over the heartbeat duration the reconstructed displacement field **d**^*n*^(**x**) (Sect. 2.3). Instead, the PV motion is achieved by prescribing opening and closing intervals by images (Fumagalli et al., 2020). The valve dynamics is handled by a cosine interpolation between the fully closed and fully open valve configurations.

#### 2.5.2 Details on the meshes

As illustrated in Fig. 3, we build tetrahedral meshes of the RH cavity with heterogeneous size *h* for the healthy (H) and for the repaired-ToF (ToF) cases, exploiting the vmtk meshing tools. The ToF mesh is used in both the PVReg-ToF and PVnoReg-ToF scenarios. We refine further the mesh element size in the regions occupied by TV and PV during their motion. This refinement is done up to half the leaflets’ thickness (we assume a constant thickness throughout the leaflets equal to 1.5*mm*) in order to correctly represent the valves using the RIIS method. Notice that we do not need to build any mesh for the leaflets’ valves as they are treated as immersed surfaces according to the RIIS method (Fedele et al., 2017). Details about the constructed meshes are reported in Table 1.

**Table 1.** Details about the tetrahedral meshes used for the DIB-CFD simulations. Total DOFs include both pressure and velocity DOFs and refer to linear finite elements.

| Patient | H | PVReg-ToF/PVnoReg-ToF |
| --- | --- | --- |
| Number of cells | 3,927,369 | 2,169,648 |
| Total DOFs | 2,578,280 | 1,462,132 |
| $h_k^{max}$ [mm] | 3.83 | 3.88 |
| $h_k^{mean}$ [mm] | 0.85 | 0.99 |
| $h_k^{min}$ [mm] | 0.32 | 0.32 |

#### 2.5.3 Setup of numerical simulation

From the numerical standpoint, we discretize the DIB-CFD problem in space by adopting inf-sup stabilized P1 − P1 finite elements and a SUPG stabilization for advection-dominated flows (Tezduyar and Senga, 2006). In time, we employ a semi-implicit backward Euler scheme with Δ*t* = 10^−5^*s*. This time step and the mesh size (reported in Table 1) are selected upon conducting a convergence test as in Bennati et al. (2023a): finer meshes or smaller time steps do not yield significant difference in the results.

We simulated 7 heartbeats with period *T* equal to 0.91*s* and 0.96*s* for H and ToF, respectively, starting from a null initial condition. Finally, we prescribe pressure curves, which are reported in Fig. 3, as boundary conditions at the inlets Σ_in_ and the outlet Σ_out_. Specifically, physiological pressures for H scenario are taken from Pappano and Wier (2018), while pathological pressure curves for PVReg-ToF and PVnoReg-ToF scenarios are obtained from a calibrated lumped parameter model (details can be found in Criseo et al. (2024)).

We solve numerically the fluid dynamics problem in the moving domains described in Sect. 2.5.1 and simulate: the healthy (H), the regurgitant repaired-ToF (PVReg-ToF), and the post-operative non-regurgitant repaired-ToF (PVnoReg-ToF) scenarios. The simulations are executed in life^x^ (Africa, 2022; Africa et al., 2024) (Finite Element, FE, C++ library based on deal.II Arndt et al. (2022) FE core and developed at MOx, Politecnico di Milano, within the framework of the iHEART project https://iheart.polimi.it) and carried out using 144 parallel processes on the Tier-0 EuroHPC super-computer LEONARDO (hosted by Cineca high-performance computing center https://www.hpc.cineca.it,Bologna, Italy).

### 2.6 Computation of postprocessed quantities

In order to remove the unphysical influence of the initial null solution, we compute the quantity of interest discarding the first heartbeat. To analyse the overall topology of the flow field, we consider the ensemble blood velocity defined as follows:

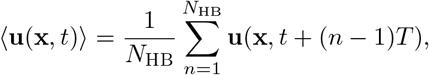

where **u** is the instantaneous blood velocity and *N*_HB_ refers to the total number of simulated heart-beats discarding the first one. We indicate with the symbol ⟨·⟩ the average over *N*_HB_ heartbeats of period *T*. We compute the velocity fluctuation field **u**^*′*^ by adopting the Reynolds decomposition (Pope, 2001) **u** = ⟨**u**⟩ − **u**^*′*^, and we analyse:

- The Reynolds number (Yoganathan et al., 1988) across TV and PV, computed as:

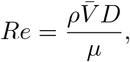

where *ρ* and *µ* are the blood density and molecular viscosity, respectively. We indicated with *D* the diameter of the equivalent circle area of the valve orifice, and with 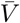 the cross sectional fluid velocity obtained by spatially averaging the total flow over the valve orifice area;
- The root mean square and the standard deviation of the velocity fluctuations indicated as 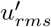 and ***σ***, respectively, and defined as (Pope, 2001):

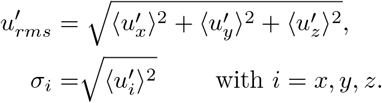

These allow us to quantify and localize the velocity fluctuations among the heartbeats;
- The total mean kinetic energy (MKE) and the total turbulent kinetic energy:

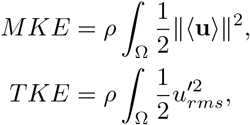

where, in particular, the TKE allows to identify the temporal instants where velocity fluctuations are more pronounced (Pope, 2001):
- The Turbulence Intensity (TI) (Yoganathan et al., 1988; Pope, 2001):

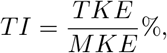

which measures the global strength of turbulence in the flow.

Finally, to quantify the stresses on the wall exerted by the blood flow and the possible presence of blood stagnation regions near the endocardium, we compute the wall shear stress (WSS), the time average wall shear stress (TAWSS) and the relative residence time (RRT) (Himburg et al., 2004; Sugiyama et al., 2013; Riccardello Jr et al., 2018; Bennati et al., 2023a).

## 3 Reconstruction results

In this section, we focus on the reconstruction results obtained by exploiting the MSMorph technique (Renzi et al., 2023) and the procedures described in Sect. 2.3 and 2.4 for the RH and the TV and PV valves, respectively.

In Table 2, we report the indexed RV and RA volumes. Differences in terms of morphology can be qualitatively appreciated in Fig. 4, left column, which depicts H and PVReg-ToF/PVnoReg-ToF’s RH in the reference end-systolic configuration. Regarding wall dynamics, we observe a relevant displacement of the intraventricular septum at end diastole in PVReg-ToF/PVnoReg-ToF as compared to the control (see Fig.4 middle column). Further alterations of the PVReg-ToF/PVnoReg-ToF RH dynamics are revealed by the atrial and ventricular volume trends reported in Fig. 4, left column. Specifically, the PVReg-ToF/PVnoReg-ToF RA is characterized by a poor contraction and its contribution to the ventricular diastolic filling due to the atrial kick is almost absent. As a consequence, we observe a prolonged ventricular diastasis.

**Table 2.** Patients’ volumetric indexes. BSA = body surface area; RVEDVi = indexed right ventricle end-diastolic volume; RVESVi = indexed right ventricle end-systolic volume; RAEDVi = indexed right atrium end-diastolic volume; RAESVi = indexed right atrium end-systolic volume.

| Patient | BSA [$m^2$] | RVEDVi [$ml/m^2$] | RVESVi [$ml/m^2$] |
| --- | --- | --- | --- |
| H | 1.96 | 100.6 | 59.2 |
| PVReg-ToF/<br>PVnoReg-ToF | 1.25 | 178.93 | 113.5 |
| | | RAEDVi [$ml/m^2$] | RAESVi [$ml/m^2$] |
|  |  | 37.2 | 61.0 |
|  |  | 52.2 | 63.6 |

**Fig 4.**
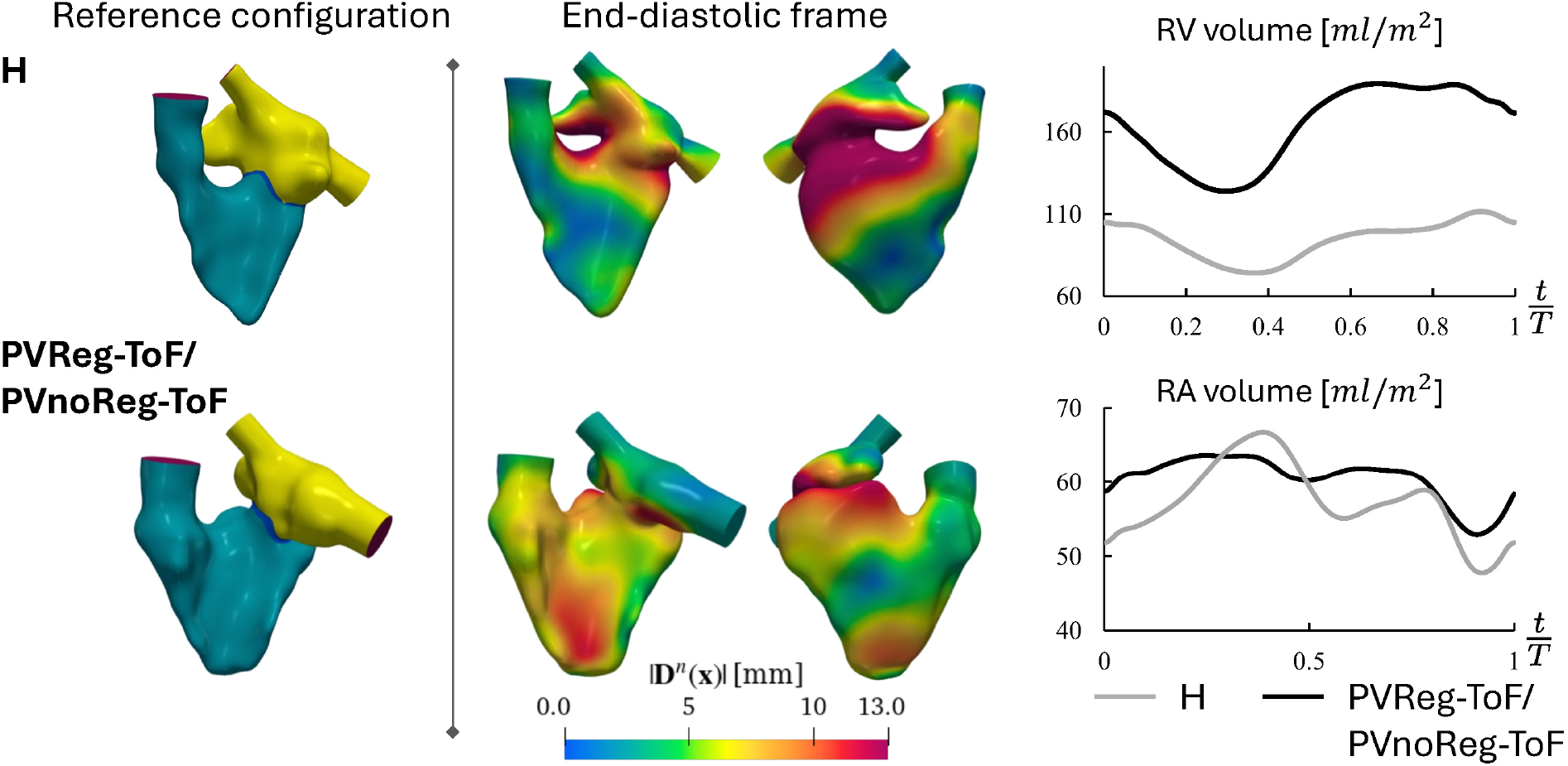
Reconstructions of the RH in the reference configuration and displacement fields for the healthy (H) and rToF (ToF) subjects. On the left column, reference configurations at end-systole with region tags: yellow -right atrium; blue -TV annulus; turquoise green -right ventricle; red -inflow and outflow sections. On the middle column, magnitude of **D**^*n*^ at end-diastole in the corresponding geometrical configuration. On the right column, ventricular and atrial volume curve over time for both subjects. Times are normalized w.r.t. the subject’s heartbeat period.

Concerning valves, we depict in Fig. 3 – b the reconstructed TV for both subjects in the fully closed and open configurations. In order to qualitatively assess the fidelity to the image, Fig. 5 presents the superposition of these results on 4 chambers and right ventricle outflow tract (RVOT) cine-MRI planes. Values of TV Effective Orifice Area (EOA) at the diastolic E-wave and A-wave instants are provided in Table 3. These quantities are calculated in post-processing as the area enclosed by the valve at the plane of the leaflet free margins.

**Table 3.** Characterization of Tricuspid (TV) and Pulmonary (PV) valves. The TV effective orifice area at the E-wave and A-wave instants are indicated by *EOA*_Ew_ and *EOA*_Aw_, respectively; *PRA* indicates the Pulmonary Regurgitant Area; *t*_*o*_*/T* and *t*_*c*_*/T* correspond to the normalized times of the onset of TV opening and closing (PV closing and opening, respectively), respectively, expressed as a fraction of the subject’s heartbeat duration.

| Patient | $EOA_{Ew} [cm^2]$ | $EOA_{Aw} [cm^2]$ | $PRA [cm^2]$ | $t_o/T$ | $t_c/T$ |
| --- | --- | --- | --- | --- | --- |
| H | 10.1 | 6.35 | — | 0.36 | 0.92 |
| PVReg-ToF/PVnoReg-ToF | 4.36 | 3.12 | 5.84 | 0.34 | 0.87 |

**Fig 5.**
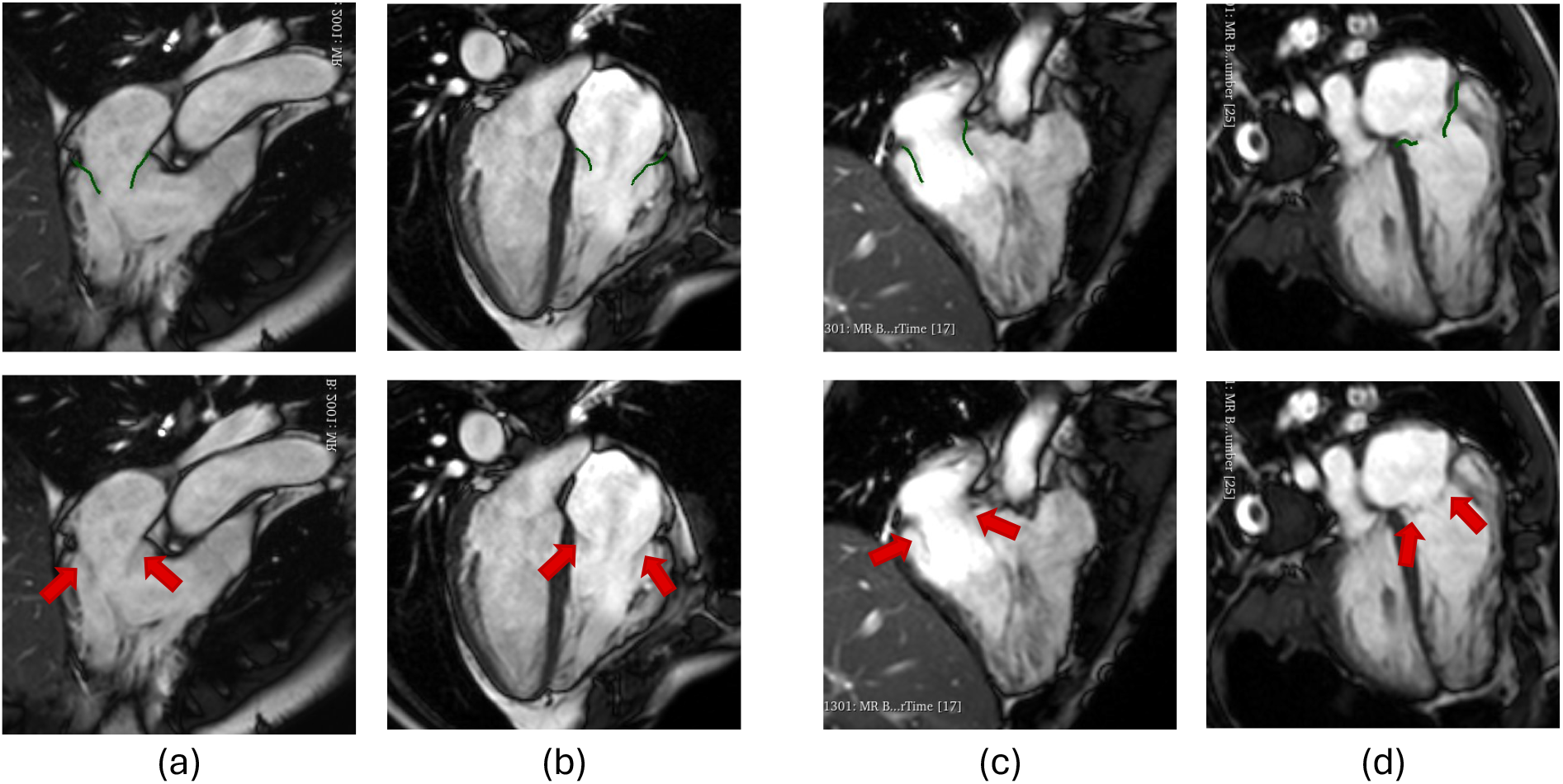
Top row: intersection of the reconstructed TV surfaces (green line) with the cine-MRI planes for the healthy subject (columns a and b) and the repaired-ToF patient (columns c and d); bottom row: same images without reconstructed surfaces, red arrows indicate the patient’s leaflets. Columns a and c: 4-chamber plane; columns b and d: RVOT plane Naoum et al. (2017).

The results of adapting the *Zygote* template PV to the subjects’ annulus are showed in Fig. 3 -b for both the PVReg-ToF and PVnoReg-ToF scenarios. In particular, we recall that to simulate the patient regurgitant fraction (RF), in the PVReg-ToF scenario, we further modify the PV template by leaving a gap between the leaflets free margins (see Sect. 2.4). Specifically, to replicate a *RF* = 38% we impose a partial closure of 50%, which lead to a pulmonary regurgintant area (PRA) of about 5.84*cm*^2^ (see Table 3). In addition, Table 3 reports the normalized times indicating the onset of TV opening and closing (PV opening and closing, respectively). Fully TV closure is reached at the end of the heartbeat for both subjects.

## 4 Numerical results

In this section, we present the results of the DIB-CFD analysis in the three scenarios presented in Sect. 1, namely, a healthy control (H), a repaired-ToF with severe pulmonary regurgitation (PVReg-ToF), and a post-operative non-regurgitant repaired-ToF (PVnoReg-ToF). The numerical results are organized as follows: in Sect. 4.1, we describe the flow topology by analysing the ensemble velocity field; in Sect. 4.2, we address the analysis of the transition to turbulence; in Sect. 4.3, we analyse stresses on walls and indicators of possible blood stagnation.

### 4.1 Characterization of flow field patterns

In this section, we report and describe the ensemble velocity field ⟨**u**⟩, the flows across the pulmonary and tricuspid valves, and the global mean kinetic energy (MKE) for the RA and RV.

The maximum ensemble velocities and flows through valves are reported in Table 4 for the three scenarios at three relevant instants: peak systole and peak diastolic E-wave and A-wave. In all three cases, the pulmonary jet during systole was characterized by maximum velocity values at the pulmonary valve orifice comparable to physiological 4D flow measurements available in literature (Hudani et al., 2023; François et al., 2012). In contrast, only for H, the blood velocities through TV are within the physiological ranges (< 0.7*m/s*), while PVReg-ToF and PVnoReg-ToF show a stenotic behaviour (Baumgartner et al., 2009). Fig. 6 illustrates the time trends of pulmonary and tricuspid flows throughout the cardiac cycle together with 3D representations of the ensemble velocity magnitude. Notably, in the H subject, we observed that the flow entering the RV through TV during the early diastole did not extend into the apical region. This behaviour was further highlighted by the flow streamlines depicted in Fig. 7. In addition, we detected a separation among the inward flows from the inferior vena cava (IVC) and the superior vena cava (SVC), persisting throughout most of the diastole. Specifically, the blood coming from IVC impinged on the RV anterior wall and then flowed mainly towards the RVOT. At the same time, blood flow from SVC moved deeper towards the ventricular apex without reaching it and drifted upwards behind the tricuspid leaflets. In the PVReg-ToF and PVnoReg-ToF cases, at the peak E-wave instant, a similar division of the flows from SVC and IVC was present (see arrows in Fig. 7).

**Table 4.** Magnitude of the ensemble velocity and values of maximum flows at three instants: *t*_*ps*_ indicates peak systole; *t*_*Ew*_ and *t*_*Aw*_ indicate peak E-wave and A-wave, respectively; TVR indicates the Tricuspid Regurgitant Volume. Flow values are normalized w.r.t the patients’ BSA.

| Scenario | $\ \langle \mathbf{u} \rangle\ _{t_{ps}} [m/s]$ | $\ \langle \mathbf{u} \rangle\ _{t_{Ew}} [m/s]$ | $\ \langle \mathbf{u} \rangle\ _{t_{Aw}} [m/s]$ | $TRV [ml/m^2]$ |
| --- | --- | --- | --- | --- |
| H | 0.97 | 0.4 | 0.43 | 5.85 |
| PVReg-ToF | 1.1 | 1.0 | 0.68 | 5.92 |
| PVnoReg-ToF | 1.3 | 1.9 | 0.7 | 16.9 |
| | $Q_{t_{ps}}^{PV} [ml/s \cdot m^2]$ | $Q_{t_{Ew}}^{TV} [ml/s \cdot m^2]$ | $Q_{t_{Aw}}^{TV} [ml/s \cdot m^2]$ | |
|  | 156.7 | 136.4 | 97.0 |  |
|  | 248.7 | 242.8 | 54.6 |  |
|  | 248.17 | 378.8 | 46.0 |  |

**Fig 6.**
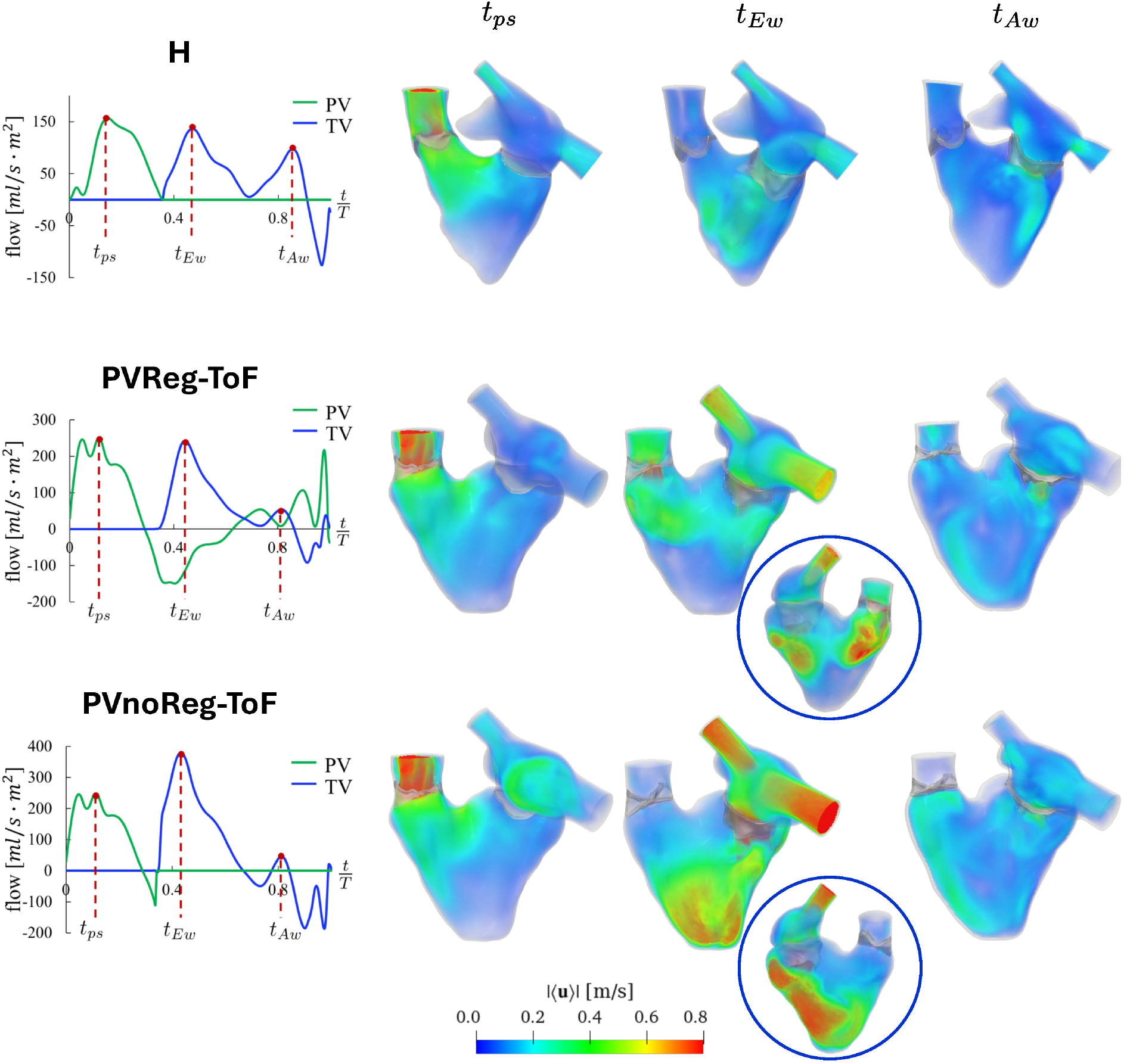
Valves’ flows and ensemble velocity magnitude for the three simulated scenarios: the healthy control (H), the repaired-ToF with severe pulmonary regurgitantion (PVReg-ToF), and the post-operative non-regurgitant repaired-ToF (PVnoReg-ToF). On the left columns, trends of flows through the pulmonary (PV) and the tricuspid valve (TV), with indication of three relevant frames: *ps* indicate the peak systole; *Ew* indicates the peak E-wave in early diastole; *Aw* indicates the peak A-wave in late diastole. On the right, the volume rendering of the ensemble velocity magnitude at the same instants. Blue circles: RH anterior view focusing on the jet impingement regions.

**Fig 7.**
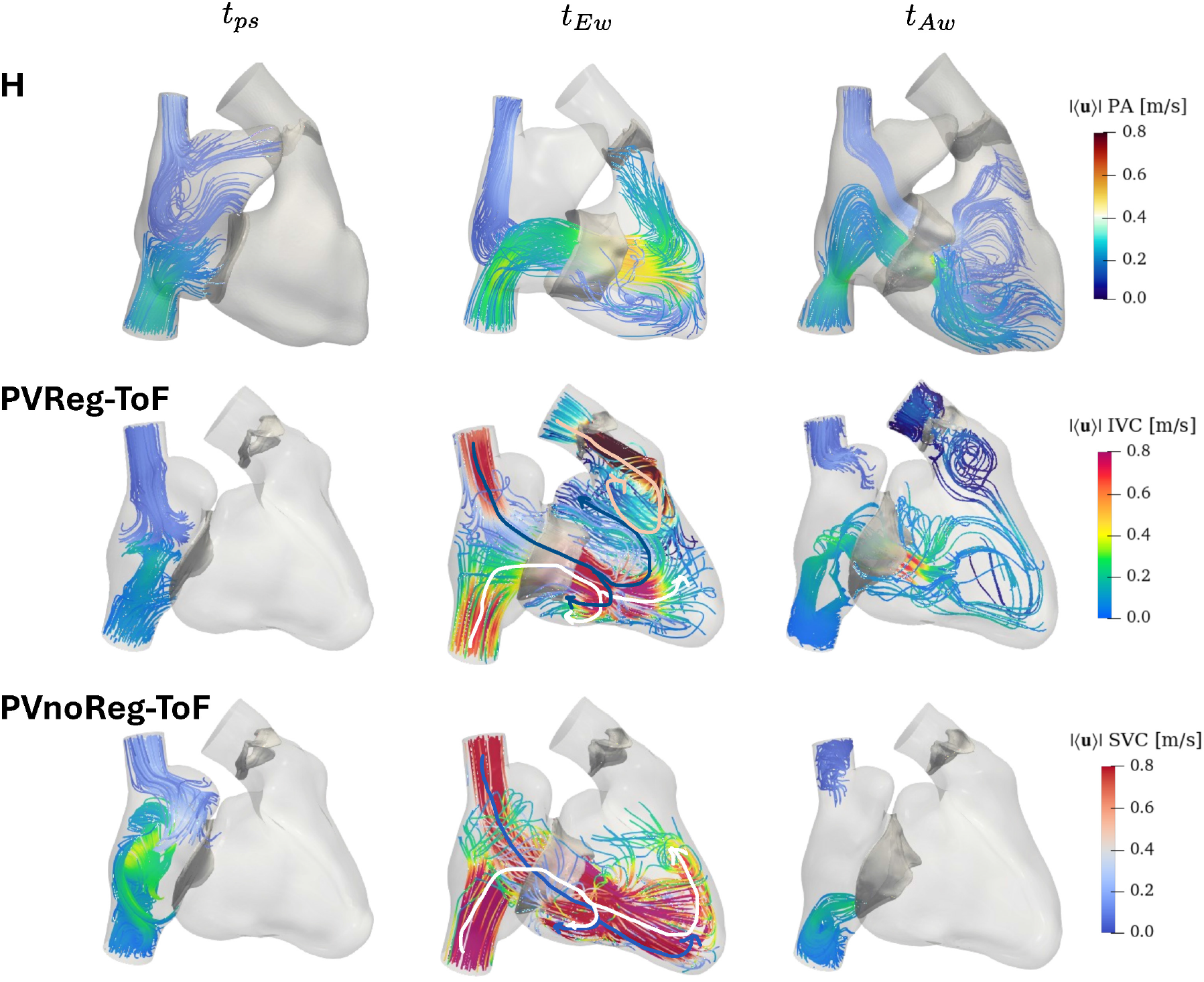
Flow streamlines at three instants (peak systole *t*_*ps*_, peak E-wave *t*_*Ew*_, and peak A-wave *t*_*Aw*_) for the three simulated scenarios: the healthy control (H), the repaired-ToF with severe pulmonary regurgitation (PVReg-ToF), and the post-operative non-regurgitant repaired-ToF (PVnoReg-ToF). Different colour bars are used to indicate flows from the pulmonary artery (PA), inferior vena cava (IVC), and superior vena cava (SVC). Arrows indicate the main direction of flows: pink for the blood flow from PA; blue for the blood flow from SVC; white for the blood flow from IVC.

However, in the PVReg-ToF scenarios, the flow from IVC reached the ventricular apex, while the blood from SVC, after impinging on the anterior wall, deflected upwards. Both PVReg-ToF and PVnoReg-ToF exhibited high velocities within the tricuspid orifice in diastole. Moreover, TV leaflets oriented the high-velocity tricuspid jet towards a region of the anterior wall closer to the posterior wall.

Concerning the pulmonary regurgitation in the PVReg-ToF scenario, a counter-rotatory flow coming from the pulmonary artery was present and acted as a barrier that prevented the tricuspid flow from reaching the RVOT (see pink arrow in Fig. 7). The peak regurgitation occurred at 40% of the heartbeat and was characterized by a maximum flow of 149.6*ml/s* · *m*^2^ with a maximum velocity of 1.7*m/s*. High regurgitant blood velocities persisted at the peak E-wave instant, about 1.3*m/s*, and inverted direction only during the late diastole, with blood velocity approximately 0.18*m/s* at peak A-wave. Also, the RV flow streamlines became more disorganized in this phase (see Fig. 7).

In the PVnoReg-ToF case, given that it shared the same stroke volume as PVReg-ToF, the presence of a competent PV caused a relevant increase of tricuspid flow in diastole, see Tab. 4. Additionally, the tricuspid jet impingement region on the anterior wall extended to the ventricular apex (see Fig. 6) and, in the absence of a re-entering pulmonary flow, the tricuspid flow was able to directly reach the RVOT (see white arrow in Fig. 7).

The RV and RA MKE time evolutions are reported in Fig. 8. For the RV, the typical three-peak curve (Barker et al., 2019) characterized all scenarios. Regarding the RA MKE, we observed that the peak during early diastole is more pronounced in PVReg-ToF and PVnoReg-ToF. This is due to the higher blood velocity through the TV in those cases. Finally, PVnoReg-ToF showed a spike in MKE at the end of diastole, attributable to the high-velocity regurgitant flow entering TV during its closure.

**Fig 8.**
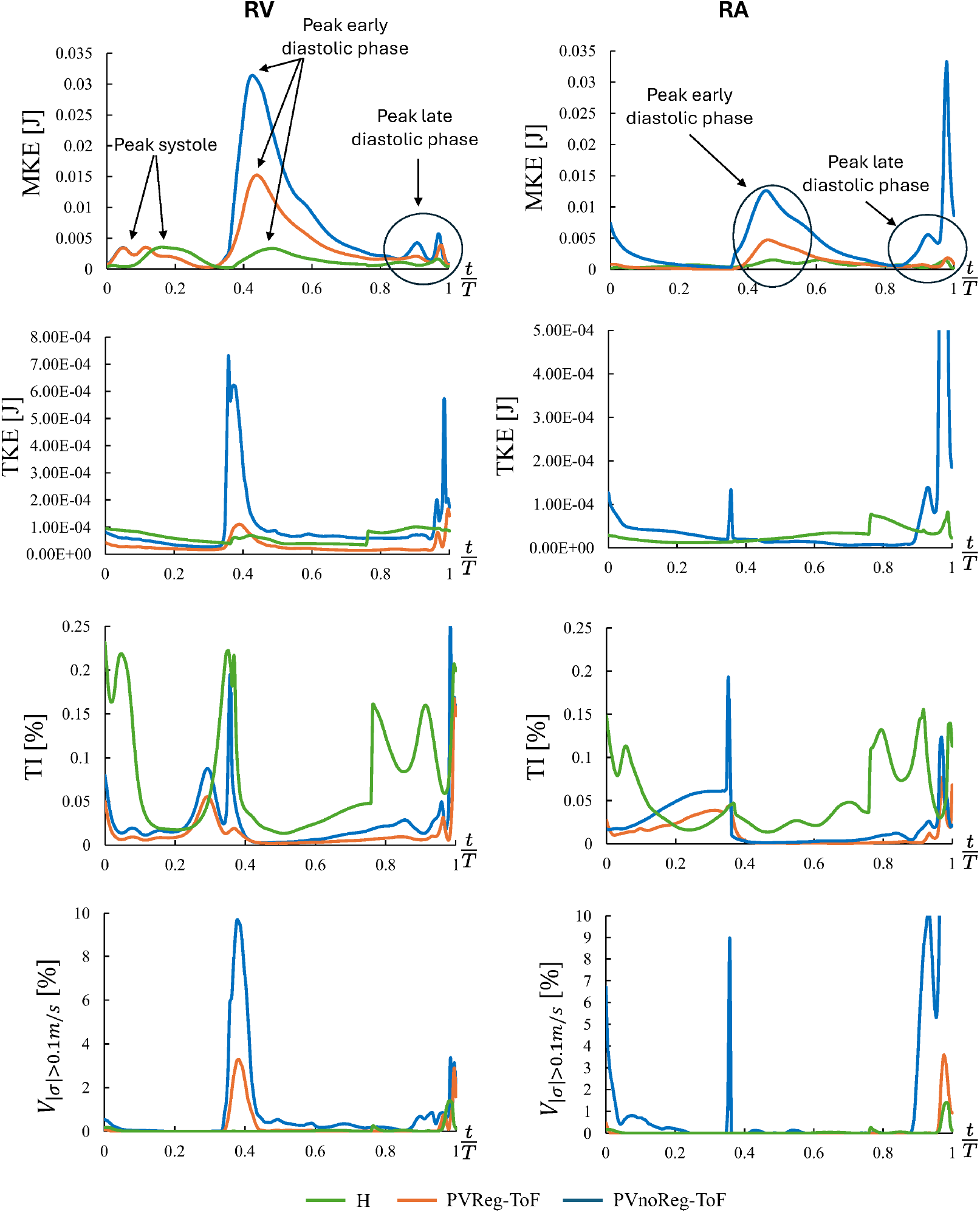
First row: global mean kinetic energy; second row: global turbulent kinetic energy; third row: turbulence intensity; fourth row: volume percentage with standard deviation of the velocity fluctuations greater than 0.1m/s. RV = right ventricle; RA = right atrium; H = control scenario; PVReg-ToF = regurgitant repaired-ToF scenario; PVnoReg-ToF = post-operative non-regurgitant repaired-ToF scenario.

### 4.2 Analysis of transition to turbulence

For the analysis of the transition to turbulence within the RH, we reported the following quantities, which are defined in Sec 2.6: in Table 5, the Reynolds numbers of the flows across PV and TV; in Fig. 9, the velocity fluctuation field 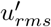 and the standard deviation of the velocity fluctuations ***σ***; in Fig. 10, the ratio between the sub-grid scale and the molecular viscosities, *µ*_*sgs*_*/µ*, and the vorticity isosurfaces; in Fig. 8, the time trends of the global turbulent kinetic energy (TKE) and of the turbulence intensity (TI). In Fig. 10, we also show the percentage of volume subjected by |***σ*** |*>* 0.1*m/s* over the cardiac cycle.

**Table 5.**
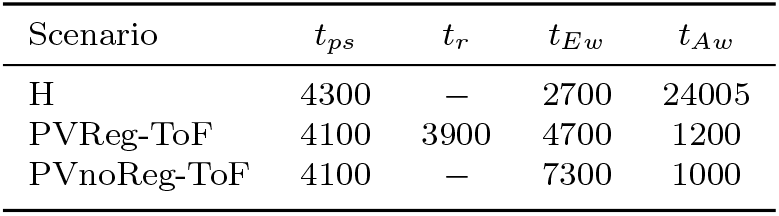
Reynolds numbers (Re) of tricuspid and pulmonary flows at four instant: *t*_*ps*_ represents peak systole; *t*_*r*_ is the instant of maximum pulmonary regurgitation; *t*_*Ew*_ represents peak E-wave; *t*_*Aw*_ represents peak A-wave

**Fig 9.**
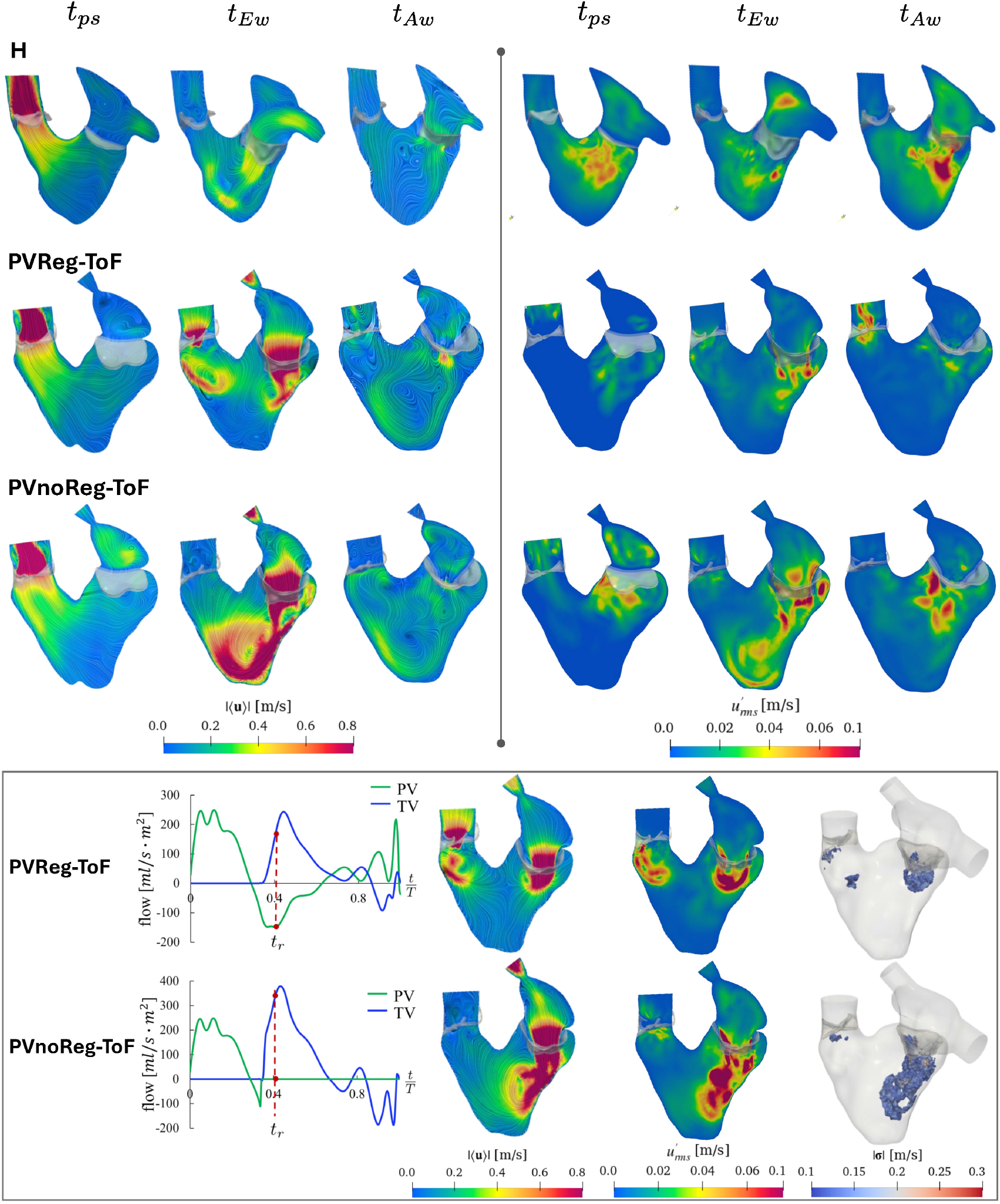
Visualization of quantities related to transition to turbulence. On top results are showed for all the scenarios at three relevant instants (peak systole, peak E-wave and peak A-wave): on the left, slices of the ensemble velocity magnitude are reported for sake of comparison; on the right, slices of the root mean square of the velocity fluctuations. Bottom panel, we report the results obtained at the instant of peak pulmonary regurgitation (*t*_*r*_) for the regurgitant (PVReg-ToF) and non-regurgitant (PVnoReg-ToF) repaired-ToF cases. From left to right: valve flows, magnitude of ensemble velocity, root mean square of the velocity fluctuations, and isosurfaces of standard deviation of the velocity fluctuations ∥***σ***∥ *>* 0.1*m/s*.

**Fig 10.**
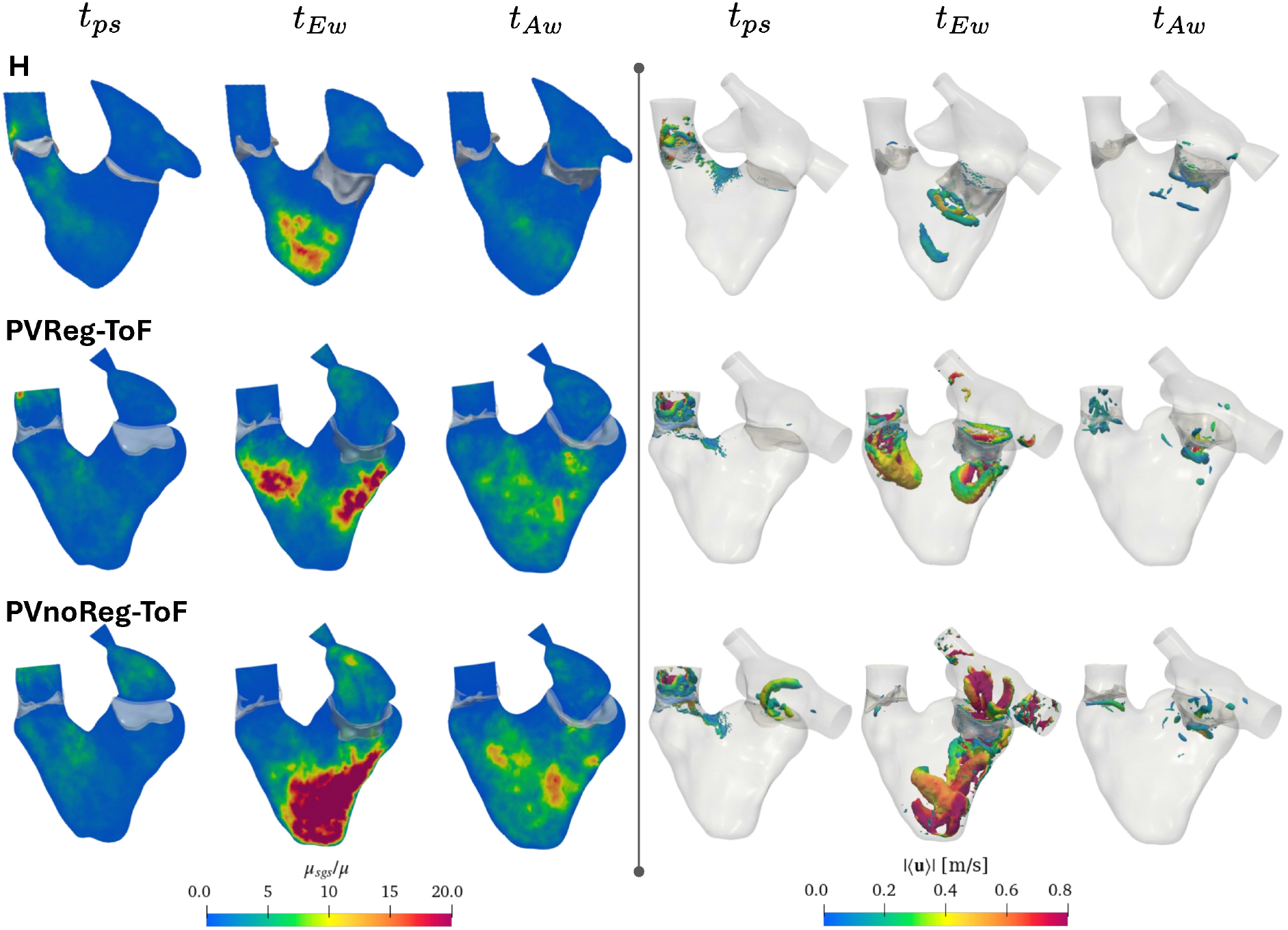
On the left, the ratio between the subgrid-scale viscosity and the molecular viscosity. On the right, vorticity isosurfaces representing the intracardiac flow, created with Q Criterion *>* 2000.

Low-velocity oscillations characterized the flow in the core of the systolic and diastolic jets, but high 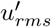 were found in the shear layers and the proximity of the valves. The highest values (about 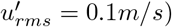 were reached at the peak E-wave within the shear layers of the tricuspid jets in the H scenario. In PVReg-ToF and PVnoReg-ToF cases, velocity oscillations had the same order of magnitude as the ensemble velocity, particularly at peak E-wave. Nonetheless, the highest values were recorded at the instant of maximum regurgitant pulmonary flow and were about 0.29*m/s* and 0.63*m/s* for PVReg-ToF and PVnoReg-ToF, respectively. Regarding the standard deviation, ***σ***, its patterns resembled those of 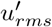 in all scenarios. The peak values of ***σ*** were identified during late diastole, at the peak A-wave, in the cases of H and PVnoReg-ToF, and corresponded approximately to 40% and 23% of the ensemble velocity, respectively. For PVReg-ToF, the peak values occurred during the early diastole, at the peak E-wave, and were around 20% of the ensemble velocity. Moreover, we identified the regions with high oscillatory behaviour, ∥***σ***∥ *>* 0.1*m/s*, in the shear layers surrounding the tricuspid and regurgitant jets (see Fig. 9). These flow regions are visualised in Fig. 9. Values of *µ*_*sgs*_*/µ* displayed in Fig. 10 proved that the sigma-LES turbulence model is active in all three scenarios, especially within the RV in diastole. The *µ*_*sgs*_ reached the highest values during E-wave: about ten times greater than *µ* in H and more than 20 times in PVReg-ToF and PVnoReg-ToF. Looking at the isosurfaces displayed in Fig. 10, ring vortices detaching from the PV leaflets were detected at peak systole. These vortices tended to rise along the pulmonary artery and, in the control case, they dissipated before reaching the outlet section, which is just before the pulmonary bifurcation plane. Additionally, below the TV, the typical “doughnut”-shaped vortex ring (ElBaz et al., 2014; Loke et al., 2022) developed in the H case at peak E-wave, and it dissipated before reaching the ventricular apex. In PVReg-ToF case, the vortices originating from the two valves, though more extensive, also dissipated before reaching the RV apex. In contrast, in the PVnoReg-ToF scenario, the vorticose structure surrounding the tricuspid high-velocity jet extended all the way to the apex. Moreover, this case presented additional vortices within RA in systole and during the early diastole, which were consistent with flow streamlines depicted in Fig. 7. Despite these findings, the recorded TI remained below 0.3% and comparable with H both within RA and RV (see Fig. 8).

### 4.3 Analysis of wall stresses and stagnation risk

Wall shear stresses (WSS) for the three scenarios are illustrated in Fig. 11. Overall, WSS remained below 1*Pa* throughout most of the heartbeat. However, peaks exceeding 2*Pa* were recorded in systole near the pulmonary leaflets’ commissures and at the supraventricular crest in all scenarios. At peak E-wave in PVReg-ToF and PVnoReg-ToF cases, elevated stresses (about 2*Pa*) were found in the anterior wall region where the high-velocity tricuspid jet impacts, as well as on the vascular walls of SVC and IVC due to increased inflow velocity. TAWSS results further proved that high WSS were transient. The highest TAWSS values stayed around 1*Pa*, and it was observed on the RVOT wall near the PV annulus, on the pulmonary artery wall, and exclusively in the PVnoReg-ToF case, on the septal wall near TV. The regions where the flow was slow (∥⟨**u**⟩∥ < 0.2*m/s*), such as at the level of the ventricular apex, within RA, and within flow shear layers, exhibited high RRT values. In particular, *RRT >* 10*Pa*^−1^ was recorded at the apex and on RA wall in the H and PVReg-ToF scenarios. Conversely, in the PVnoReg-ToF case, the high-velocity tricuspid, SVC, and IVC flows effectively washed out these regions.

**Fig 11.**
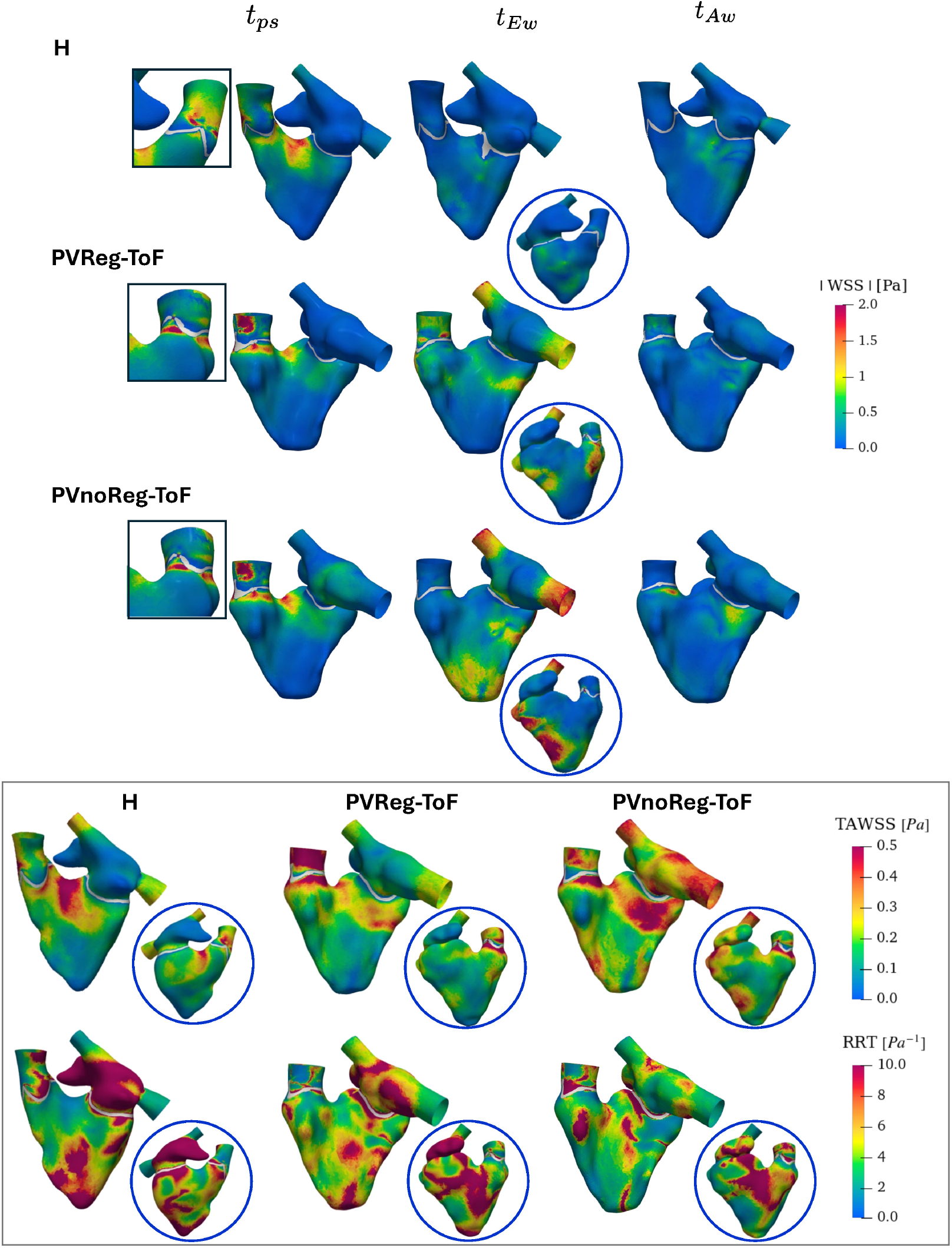
Stresses and stagnation risk. On top, a representation of the 3D wall shear stress (WSS) patterns on the endocardial walls; square boxes: focus on the pulmonary commissures; round boxes: visualization of the anterior wall. Bottom panel: time averaged WSS (TAWSS) and relative residence time (RRT) patterns on the first and second rows, respectively, for the three scenarios.

## 5 Discussion

We structure the discussion into distinct points to emphasize key aspects of the right heart hemodynamics that we have identified. Specifically, we highlight: the novelties with respect to previous studies (Sect. 5.1); the specific behaviours of the RV hemodynamics with respect to the LV one (Sect. 5.2); the differences between healthy and repaired-ToF cases (Sect. 5.3); the characterization of turbulence effects (Sect. 5.4); the influence of pulmonary valve replacement (Sect. 5.5).

### 5.1 Novelties of the present work

Characterizing flow within the right heart (RH) is particularly challenging due to its complex geometries. However, gaining insight into RH fluid dynamics is of extreme diagnostic importance because of the high incidence of acquired and congenital heart diseases with impaired RH functionality (Dumitrescu et al., 2018; Valsangiacomo Buechel and Mertens, 2012; Haddad et al., 2008). With this aim, several studies have employed 4D flow MRI to investigate the intracardiac flow patterns within RH (François et al., 2012; Hudani et al., 2023; Barker et al., 2019; ElBaz et al., 2014; Hirtler et al., 2016).

In this work, we carried out a computational study of the fluid dynamics of the RH. Our computational models were fully patient-specific and incorporated both chambers and valves with their motion. To the best of the authors’ knowledge, this is the first work that addresses the analysis of fluid dynamics within a complete patient-specific RH model (RV + RA + TV + PV + PA) via DIB-CFD simulations. Previous computational studies on the RH focused on either the ventricular (Mangual et al., 2012; Wiputra et al., 2018; Loke et al., 2022, 2024) or the atrial chamber (Mareels et al., 2004; de Oliveira et al., 2021; Parker et al., 2022), separately. Knowing that RH efficiency depends on the RA-RV interplay, we considered the inclusion of both chambers crucial, particularly in the context of ToF, where it has been proved that the RA performance is an independent predictor for adverse outcomes post-PVR (Ali et al., 2019; Ait-ALi et al., 2020).

Moreover, as far as we are aware, this is the first computational study that includes the patient’s TV reconstructed by images; prior works on RH hemodynamics typically modelled it by a planar surface (Loke et al., 2022; Wiputra et al., 2018).

Finally, we introduced a novel method that leverages information from all the available series to reconstruct the TV shape and motion from multiple cine-MRI acquisitions. This allowed us to obtain patient-specific TV without relying on specific acquisitions, such as rotational cine-MRI or three-dimensional echocardiography. Indeed, the proposed technique allowed for the first time, to obtain it by using only long-axis and 2/3/4 Chambers views. Owing to this, we were able, for the first time, to simulate the presence of mild tricuspid regurgitation both in healthy and pathological patient-specific scenarios, as well as the stenotic behaviour of the repaired-ToF TV.

Notably, the proposed procedure can also be exploited for other dynamic images besides cine-MRI, such as echocardiography, due to its flexibility with respect to the available medical images.

### 5.2 Comparison between right and left ventricular hemodynamics

Our investigation highlighted key differences and commonalities between the RV and LV hemodynamics.

Similarly to the LV, the healthy RV generated a ring-shaped vortex during early diastole, which rolled up at the free margins of the TV leaflets. However, unlike in the LV, this RV vortex dissipates before reaching the apical region, mainly due to the lower inertia of the tricuspid jet (ElBaz et al., 2014; Seo et al., 2014; Collia et al., 2021).

Despite these differences, the intraventricular hemodynamics is macroscopically similar between the left and right sides. As illustrated in Fig. 12, the inflow develops into two counter-rotating flows in both ventricles: a dominant one directed towards the outflow tract and a secondary, smaller one behind the valve leaflets. This flow pattern ensures that blood efficiently reaches the outflow and enters the circulatory system. However, these processes occur in two completely morphologically different chambers: an ellipsoidal-like shape in LV and a “crescent” shape in RV. Consequently, the atrioventricular valve play a crucial role in directing blood towards the semilunar valves, with their anatomy and morphology that are tailor-made to account for the different chambers’ shapes.

**Fig 12.**
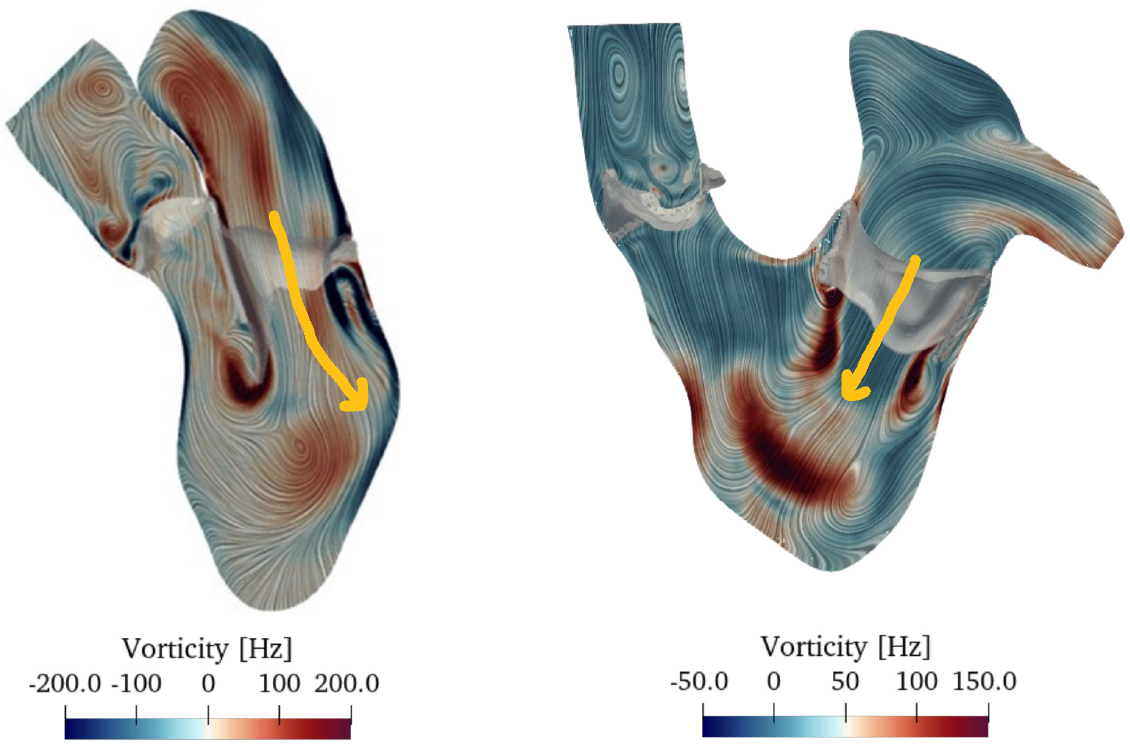
Spatial distribution of the vorticity field at the peak of E-wave for two representative healthy cases: left heart of and aorta on the left; right heart and pulmonary artery on the right. Notice, highlighted by the yellow arrows, the straight inflow through the atrioventricular valve in the right ventricle, in contrast to the deflected blood flow entering the left ventricle, as a consequence of the asymmetric atrioventricular (mitral) valve. The latter picture has been modified from the PhD thesis by Lorenzo Bennati, University of Verona.

Specifically, in RV, the chamber’s curvature (in particular, at the level of the septum and of the crista terminalis) facilitates the blood flow into PA, after it impinges on the anterior wall; thus, a standard symmetric valve with three homogeneous leaflets is sufficient for ensuring the effective RV ejection. In contrast, the LV’s non-standard asymmetric valve with two completely different leaflets allows to avoid blood stagnation in the apical region. Indeed, it redirects the mitral jet to create a circulatory flow toward the aortic valve.

Furthermore, we observe that the presence of a second counter-rotating flow within the RV facilitates blood mixing of the apical blood particles, which are not reached by the main flow.

Finally, for sake of comparison with the LV literature, we computed the standard deviation of the fluctuations of the velocity magnitude, *SD*, as defined in Vergara et al. (2017); Stella et al. (2019). We found that the peak value of this quantity during diastole was about 0.1 − 0.17*m/s*, corresponding to 25 − 33% of the mean velocity. Similar results were detected within the left chamber in a recent work (Bennati et al., 2023a). This may suggest that, despite the hemodynamics conditions differ among LV and RV, namely intrachamber pressures and mean blood velocities, the mechanisms that regulate the balance between inertial and viscous forces act in such a way to ensure a comparable filling efficiency in healthy conditions.

### 5.3 Characterization of the regurgitant repaired-ToF hemodynamics

Thanks to our patient-specific DIB-CFD simulations, we were able to characterize in detail the complex pattern of the velocity field in both healthy and congenital RH (H and PVReg-ToF, respectively).

The two simulated scenarios differed in terms of chambers and valves’ geometry and motion, as well as in terms of boundary conditions. These differences collectively contributed to determining their specific hemodynamics. For the PVReg-ToF case, a key distinctive feature is the development of a high-velocity jet through the TV at peak E-wave (see Fig. 6), which impinged on the RV lateral wall, causing a localized increase of WSS (see Fig. 11). The deflection of the diastolic jet towards the later ventricular wall, rather than the RVOT, partially contrasts with recent in-vitro findings on the effect of pulmonary regurgitation on the fluid dynamics in repaired-ToF RV presented in Mikhail et al. (2020). This discrepancy my be attributable to the presence of the patientspecific valve in our model, which was not included in Mikhail et al. (2020). Indeed, it is know that the evolution of flow within the chambers is deeply affected by the valve morphology (Seo et al., 2014; Bennati et al., 2023b) together with the ventricular wall dynamics. Specifically, the PVReg-ToF TV was slightly stenotic compared to the healthy one (*EOA* = 4.36*cm*^2^ against 10.1*cm*^2^ of H scenario, Table 3), leading to a pressure drop across the valve of about 7*mmHg*, which is outside the physiological ranges 1 −3*mmHg* (Baumgartner et al., 2009) (Δ*p* = 2*mmHg* for H). Additionally, the cardiac motion determined very high TV flow at peak E-wave (see Table 4).

Another interesting topic related to the repaired-ToF simulation is represented by the atrium modelling. The reconstruction results highlighted the pathological behaviour of the PVReg-ToF RA during diastole. In particular, Fig. 4 revealed the characteristic low compliance and increased conduit function that are typical features of a repaired-ToF RA (Kutty et al., 2017). By introducing the RA-RV coupling, we were able to reproduce the effects of this impaired RA function on the RV diastolic filling, which prolonged its diastasis phase. Moreover, effects were also observed on the TV closure dynamics, which lasted about 130*ms* in the PVReg-ToF scenario against the 80*ms* of H, as it can be deduced from the normalized times reported in Table 2. Embedding the patient-specific TV dynamics in the model also enabled us to capture the tricuspid regurgitation (TR) at the end of diastole (see Table 4). TR fell within the physiological ranges (Hahn et al., 2022) both in the H and PVReg-ToF cases. As far as we are aware, this is the first time that TR has been considered in CFD simulations despite its prevalence in 65 ™ 85% of the population (Arsalan et al., 2017). Given the high incidence of ToF, being able to computationally reproduce and analyse the effects of this condition on the intracardiac fluid dynamics can be a valuable tool in aiding surgical planning in view of avoiding irreversible RV damage in severe and chronic scenarios (Chorin et al., 2020).

Furthermore, the modelling of the entire RH system allowed us to detail the specific interplay between inflows from the superior and inferior vena cava (SVC and IVC, respectively), how the two sources of flow contribute to the regional ventricular filling, and how this interplay differs between healthy and pathological conditions. These results were enhanced by the velocity streamlines depicted in Fig. 7 and were qualitatively in agreement with literature findings (François et al., 2012; ElBaz et al., 2014; Barker et al., 2019; Hudani et al., 2023) for both H and PVReg-ToF cases.

### 5.4 Characterization of transitional effects

Although the Reynolds numbers computed across valves in systole and diastole are quite high (see Table 5), approaching the upper limit of the transitional range (Yoganathan et al., 1988), we did non observe the complete transition to a fully turbulent regime in all the simulated scenarios. This can be attributable to the fact that the blood acceleration phases caused by the heart pulsatility tend to stabilize and laminarize the flow within the chambers both in systole and diastole.

Despite the absence of turbulence, there are still flow instabilities within the RA and RV, as showed by the pattern of velocity fluctuations and sub-grid scale viscosity (see Fig. 9 and 10, respectively). Moreover, our highly detailed model allowed us to capture the vorticose structure that characterize the RV hemodynamics and energy exchange. The presence of these characteristic structures have been well documented in both the computational and clinical literature. Among all, we detected the main vortex that develops from TV (see Fig. 10) during diastole and that features a very regular ring-like shape in healthy RV scenario (François et al., 2012; ElBaz et al., 2014; Collia et al., 2021; Loke et al., 2024). In the PVReg-ToF case, this diastolic ring has a more elongated shape, which might be due to the higher jet velocity, the patient’s TV shape (it is known that the leaflets of the atrioventricular valves orient the flow within the ventricle (Seo et al., 2014)), and the interaction with the pulmonary regurgitation jet as it was also detected in Loke et al. (2024). The diastolic vortex dissipates in both the healthy and pathological scenarios before reaching the RV apex. However, while for H, this dissipation is due to the relatively low Reynolds number (*RE ≈* 2700) of the tricuspid jet, in the case of PVReg-ToF, it is due to the interaction with pulmonary vortices associated with the regurgitant jet.

We adopted the Large Eddy Simulation *σ* model to describe and quantify the turbulent and transitional phenomena. This model has already proven to be suitable in simulating fluids in enclosed domain and in the presence of shear layers (Chnafa et al., 2016; Vergara et al., 2017). Although previous works described and quantified the turbulent energy and energy loss in the contest of regurgitant repaired-ToF (Collia et al., 2021; Loke et al., 2022, 2024), in this study we carried out an original analysis by exploring the contribution of the different scales to the distribution of energy within the healthy and pathological RH. In particular, by examining the global quantities, we observed that the PVReg-ToF presented a lower ratio between mean and turbulent kinetic energies (turbulent intensity, TI) compared to H, despite the higher Reynold numbers of its tricuspid and pulmonary jets (see Fig. 8). Notably, the peak values of the standard deviation of the velocity fluctuations (reported in Sect. 4.2) also reflected this trend as their percentage with respect to the ensemble velocity halved in PVReg-ToF as compared to the healthy scenario. At the same time Fig. 10 showed a huge increase of the ratio between the sub-grid scale viscosity and the molecular viscosity in the PVReg-ToF scenario, which suggests that the smallest scales contribute more to the dissipation of the turbulent energy with respect to H. In summary, in PVReg-ToF case, the laminar flow predominates over turbulence on the large scale since the velocity fluctuations are dampened by the rapid transfer of the turbulent energy towards the more active smallest scales. Conversely, within the healthy RH, the turbulent energy is more homogeneously distributed among the scales.

To conclude, the observed results suggested that the chaotic phenomena at the smallest scales are more pronounced in the pathological RH, where are mainly localized below the RVOT and the TV leaflets. This also could affect the forces exerted among the fluid layers and thus potentially damage blood cells or the cardiac wall endothelium (Lu et al., 2001).

### 5.5 Fluid dynamics changes induced by pulmonary valve replacement

To investigate the hemodynamics phenomena occurring immediately after PVR intervention in repaired-ToF subject we build a non-regurgitant repaired-ToF model, PVnoReg-ToF, on top of PVReg-ToF, by replacing the regurgitant valve with a competent one and by imposing customized boundary conditions for the fluid problem. With this setting, inserting a competent PV considerably affects the overall RH hemodynamics. In particular, the most marked effects were observed in diastole, where we reported increased tricuspid flow at peak E-wave, with elevated velocities and Reynolds numbers (*Re* ≈ 7200). Moreover, the augmented kinetic energy of the jet, combined with the absence of the flow obstacle given by the pulmonary backflow (as for PVReg-ToF), increased the RV penetration depth, which extended along the entire RV longitudinal dimension. This increased load to the ventricle, in turn, led to higher WSS on the wall at the apex, though they remained transient (TAWSS values were below 0.2*Pa*) promoting, at the same time, the apical blood wash-out (see Fig. 11). Flow instabilities grew within the overall RH, especially within RA, where vortical structures appeared already in diastole (see Fig. 8 and 10). Finally, during TV closure, the TR become moderate, increasing up to 17*ml/m*^2^ (Hahn et al., 2022).

Summarizing, the findings of this pilot study suggested that PVR leads to an overall increase in the RH load. Although the presented results are still preliminary, the observed overload could trigger changes, especially in the valve dynamics and cardiac motion, which may correlate with restoring RH functionality in repaired-ToF patients.

## 6 Limitations

Some limitations characterize this work:

- We analysed only two patients. To support the considerations drawn in this work, we should consider a larger cohort of control subjects and repaired-ToF patients who underwent PVR. However, we believe that the presented results are very promising and could provide an excellent starting point for future studies;
- Due to the absence of patients’ clinical measurements, we could not validate our numerical results. Nonetheless, we demonstrated that our findings are meaningful and both qualitatively and quantitatively consistent with the literature;
- As we imposed Neumann boundary condition at the inlets of our DIB-CFD model, we could not reproduce the torsional motion of the SVC and IVC inflows (Parker et al., 2022), which could affect the blood velocity pattern and vortex formation within the right atrium;
- ToF patients are characterized by an asymmetrical flow partition among the two pulmonary arteries (Hudani et al., 2023), which could affect the downstream blood dynamics. However, we did not include the pulmonary bifurcation due to limitations of the acquired regions in the available cine-MRI data;
- We did not consider the subvalvular apparatus and trabeculations in the RH model. This is a common choice due to the challenges in accurately reconstructing the geometry and motion of these structures, which requires more spatially resolved imaging than cine-MRI. However, their presence has been demonstrated to affect the vortex organization and WSS patterns, in particular at the level of the ventricular apex (Sacco et al., 2018), potentially affecting the local blood wash-out predictions. Therefore, future studies should incorporate these structures for a more comprehensive analysis.

## Acknowledgements

FR, CV acknowledge their membership to INdAM GNCS -Gruppo Nazionale per il Calcolo Scientifico (National Group for Scientific Computing, Italy). FR has been partially supported by the INdAM GNCS project CUP E53C23001670001.

CV has been partially supported by the Italian Ministry of University and Research (MIUR) within the PRIN (Research projects of relevant national interest) MIUR PRIN22-PNRR n. P20223KSS2 “Machine learning for fluid-structure interaction in cardiovascular problems: efficient solutions, model reduction, inverse problems, and by the Italian Ministry of Health within the PNC PROGETTO HUB LIFE SCIENCE -DIAGNOSTICA AVANZATA (HLS-DA) “INNOVA”, PNC-E3-2022-23683266-CUP: D43C22004930001, within the “Piano Nazionale Complementare Ecosistema Innovativo della Salute” -Codice univoco investimento: PNC-E3-2022-23683266.

The authors acknowledge the CINECA award under the ISCRA initiative, for the availability of high performance computing resources and support.

## Declarations

### Ethical approval

Ethical review board gave approval for this study, and informed consent was obtained from all patients.

## References

Pasquale Claudio Africa. lifex: A flexible, high performance library for the numerical solution of complex finite element problems. SoftwareX, 20:101252, 2022. doi: 10.1016/j.softx.2022.101252.

Pasquale Claudio Africa, Ivan Fumagalli, Michele Bucelli, Alberto Zingaro, Marco Fedele, Luca Dede’, and Alfio Quarteroni. lifex-cfd: An opensource computational fluid dynamics solver for cardiovascular applications. Computer Physics Communications, 296:109039, 2024. ISSN 0010-4655. doi: 10.1016/j.cpc.2023.109039.

Lamia Ait-ALi, Chiara Marrone, Stefano Salvadori, Duccio Federici, Vitali Pak, Lugi Arcieri, Claudio Passino, Giuseppe Santoro, and Pierluigi Festa. Impact of right atrium dimension on adverse outcome after pulmonary valve replacement in repaired tetralogy of fallot patients. The International Journal of Cardiovascular Imaging, 36(10):1973–1982, 2020. doi: 10.1007/s10554-020-01891-9.

Lamia Ait Ali, Philipp Lurz, Andrea Ripoli, Giuseppe Rossi, Tobias Kister, Giovanni Donato Aquaro, C Passino, Philipp Bonhoeffer, and Pierluigi Festa. Implications of atrial volumes in surgical corrected tetralogy of fallot on clinical adverse events. International Journal of Cardiology, 283:107–111, 2019. doi: 10.1016/j.ijcard.2019.02.018.

Luca Antiga, Marina Piccinelli, Lorenzo Botti, Bogdan Ene-Iordache, Andrea Remuzzi, and David A Steinman. An image-based modeling framework for patient-specific computational hemodynamics. Medical & biological engineering & computing, 46:1097–1112, 2008. doi: 10.1007/s11517-008-0420-1.

Daniel Arndt, Wolfgang Bangerth, Marco Feder, Marc Fehling, Rene Gassmöller, Timo Heister, Luca Heltai, Martin Kronbichler, Matthias Maier, Peter Munch, et al. The deal. ii library, version 9.4. Journal of Numerical Mathematics, 30(3):231–246, 2022. doi: 10.1515/jnma-2022-0054.

Mani Arsalan, Thomas Walther, Robert L Smith, and Paul A Grayburn. Tricuspid regurgitation diagnosis and treatment. European heart journal, 38(9):634–638, 2017. doi: 10.1093/eurheartj/ehv487.

Christoph M Augustin, Andrew Crozier, Aurel Neic, Anton J Prassl, Elias Karabelas, Tiago Ferreira da Silva, Joao F Fernandes, Fernando Campos, Titus Kuehne, and Gernot Plank. Patient-specific modeling of left ventricular electromechanics as a driver for haemodynamic analysis. EP Europace, 18(Suppl 4): iv121–iv129, 2016. doi: 10.1093/europace/euw369.

Natasha Barker, Benjamin Fidock, Christopher S Johns, Harjinder Kaur, Gareth Archer, Smitha Rajaram, Catherine Hill, Steven Thomas, Kavitasagary Karunasaagarar, David Capener, et al. A systematic review of right ventricular diastolic assessment by 4d flow cmr. BioMed research international, 2019(1):6074984, 2019. doi: 10.1155/2019/6074984.

Helmut Baumgartner, Judy Hung, Javier Bermejo, John B Chambers, Arturo Evangelista, Brian P Griffin, Bernard Iung, Catherine M Otto, Patricia A Pellikka, and Miguel Quiñones. Echocardiographic assessment of valve stenosis: Eae/ase recommendations for clinical practice. European Journal of Echocardiography, 10(1):1–25, 2009. doi: 10.1016/j.echo.2008.11.029.

Lorenzo Bennati, Vincenzo Giambruno, Francesca Renzi, Venanzio Di Nicola, Caterina Maffeis, Giovanni Puppini, Giovanni Battista Luciani, and Christian Vergara. Turbulent blood dynamics in the left heart in the presence of mitral regurgitation: a computational study based on multi-series cine-mri. Biomechanics and Modeling in Mechanobiology, 22(6):1829–1846, 2023a. doi: 10.1007/s10237-023-01735-0.

Lorenzo Bennati, Christian Vergara, Vincenzo Giambruno, Ivan Fumagalli, Antonio Francesco Corno, Alfio Quarteroni, Giovanni Puppini, and Giovanni Battista Luciani. An image-based computational fluid dynamics study of mitral regurgitation in presence of prolapse. Cardiovascular Engineering and Technology, 14(3): 457–475, 2023b. doi: 10.1007/s13239-023-00665-3.

Giovanni Biglino, Claudio Capelli, Jan Bruse, Giorgia M Bosi, Andrew M Taylor, and Silvia Schievano. Computational modelling for congenital heart disease: how far are we from clinical translation? Heart, 103(2):98–103, 2017. doi: 10.1136/heartjnl-2016-310423.

Michele Bucelli, Alberto Zingaro, Pasquale Claudio Africa, Ivan Fumagalli, Luca Dede’, and Alfio Quarteroni. A mathematical model that integrates cardiac electrophysiology, mechanics, and fluid dynamics: Application to the human left heart. International Journal for Numerical Methods in Biomedical Engineering, 39(3): e3678, 2023. doi: 10.1002/cnm.3678.

Christophe Chnafa, Simon Mendez, and Franck Nicoud. Image-based simulations show important flow fluctuations in a normal left ventricle: what could be the implications? Annals of biomedical engineering, 44:3346–3358, 2016. doi: 10.1007/s10439-016-1614-6.

Ehud Chorin, Zach Rozenbaum, Yan Topilsky, Maayan Konigstein, Tomer Ziv-Baran, Eyal Richert, Gad Keren, and Shmuel Banai. Tricuspid regurgitation and long-term clinical outcomes. European Heart Journal-Cardiovascular Imaging, 21(2):157–165, 2020. doi: 10.1093/ehjci/jez216.

Dario Collia, Luigino Zovatto, Giovanni Tonti, and Gianni Pedrizzetti. Comparative analysis of right ventricle fluid dynamics. Frontiers in Bioengineering and Biotechnology, 9:667408, 2021. doi: 10.3389/fbioe.2021.667408.

Elisabetta Criseo, Ivan Fumagalli, Alfio Quarteroni, Stefano Maria Marianeschi, and Christian Vergara. Computational haemodynamics for pulmonary valve replacement by means of a reduced fluid-structure interaction model. International Journal for Numerical Methods in Biomedical Engineering, page e3846, 2024. doi: 10.1002/cnm.3846.

Judith AAE Cuypers, Myrthe E Menting, Elisabeth EM Konings, Petra Opić, Elisabeth MWJ Utens, Willem A Helbing, Maarten Witsenburg, Annemien E van den Bosch, Mohamed Ouhlous, Ron T van Domburg, et al. Unnatural history of tetralogy of fallot: prospective follow-up of 40 years after surgical correction. Circulation, 130 (22):1944–1953, 2014. doi: 10.1161/CIRCULATIONAHA.114.009454.

Diana C de Oliveira, David G Owen, Shuang Qian, Naomi C Green, Daniel M Espino, and Duncan ET Shepherd. Computational fluid dynamics of the right atrium: assessment of modelling criteria for the evaluation of dialysis catheters. PLoS One, 16(2):e0247438, 2021. doi: 10.1371/journal.pone.0247438.

Silviu Ionel Dumitrescu, Ion C Ţintoiu, and Malcolm John Underwood. Right Heart Pathology: From Mechanism to Management. Springer, 2018. ISBN 978-3-319-73763-8. doi: 10.1007/978-3-319-73764-5.

Mohammed S ElBaz, Emmeline Calkoen, Jos J Westenberg, Boudewijn PF Lelieveldt, Arno Roest, and Rob J van der Geest. Three dimensional right ventricular diastolic vortex rings: characterization and comparison with left ventricular diastolic vortex rings from 4d flow mri. Journal of Cardiovascular Magnetic Resonance, 16:1–3, 2014. doi: 10.1186/1532-429X-16-S1-P42.

Marco Fedele and Alfio Quarteroni. Polygonal surface processing and mesh generation tools for the numerical simulation of the cardiac function. International Journal for Numerical Methods in Biomedical Engineering, 37(4): e3435, 2021. doi: 10.1002/cnm.3435.

Marco Fedele, Elena Faggiano, Luca Dedé, and Alfio Quarteroni. A patient-specific aortic valve model based on moving resistive immersed implicit surfaces. Biomechanics and modeling in mechanobiology, 16:1779–1803, 2017. doi: 10.1007/s10237-017-0919-1.

Miguel A Fernández, Jean-Frédéric Gerbeau, and Vincent Martin. Numerical simulation of blood flows through a porous interface. ESAIM: Mathematical Modelling and Numerical Analysis, 42(6):961–990, 2008. doi: 10.1051/m2an:2008031.

Christopher J François, Shardha Srinivasan, Mark L Schiebler, Scott B Reeder, Eric Niespodzany, Benjamin R Landgraf, Oliver Wieben, and Alex Frydrychowicz. 4d cardiovascular magnetic resonance velocity mapping of alterations of right heart flow patterns and main pulmonary artery hemodynamics in tetralogy of fallot. Journal of Cardiovascular Magnetic Resonance, 14(1):11, 2012. doi: 10.1186/1532-429X-14-16.

Ivan Fumagalli, Marco Fedele, Christian Vergara, Sonia Ippolito, Francesca Nicolò, Carlo Antona, Roberto Scrofani, Alfio Quarteroni, et al. An image-based computational hemodynamics study of the systolic anterior motion of the mitral valve. Computers in Biology and Medicine, 123:103922, 2020. doi: 10.1016/j.compbiomed.2020.103922.

Ivan Fumagalli, Piermario Vitullo, Christian Vergara, Marco Fedele, Antonio F Corno, Sonia Ippolito, Roberto Scrofani, and Alfio Quarteroni. Image-based computational hemodynamics analysis of systolic obstruction in hypertrophic cardiomyopathy. Frontiers in Physiology, page 2437, 2022. doi: 10.3389/fphys.2021.787082.

Ivan Fumagalli, Stefano Pagani, Christian Vergara, Dilachew A Adebo, Maurizio Del Greco, Antonio Frontera, Giovanni Battista Luciani, Gianluca Pontone, Roberto Scrofani, Alfio Quarteroni, et al. The role of computational methods in cardiovascular medicine: a narrative review. Translational Pediatrics, 13 (1):146, 2024. doi: 10.21037%2Ftp-23-184.

Juan C Grignola. Hemodynamic assessment of pulmonary hypertension. World journal of cardiology, 3(1):10, 2011. doi: 10.4330%2Fwjc.v3.i1.10.

François Haddad, Ramona Doyle, Daniel J Murphy, and Sharon A Hunt. Right ventricular function in cardiovascular disease, part ii: pathophysiology, clinical importance, and management of right ventricular failure. Circulation, 117(13):1717–1731, 2008. doi: 10.1161/CIRCULATIONAHA.107.653584.

Rebecca T Hahn, Luigi P Badano, Philipp E Bartko, Denisa Muraru, Francesco Maisano, Jose L Zamorano, and Erwan Donal. Tricuspid regurgitation: recent advances in understanding pathophysiology, severity grading and outcome. European Heart Journal-Cardiovascular Imaging, 23(7):913–929, 2022. doi: 10.1093/ehjci/jeac009.

Heather A Himburg, Deborah M Grzybowski, Andrew L Hazel, Jeffrey A LaMack, Xue-Mei Li, and Morton H Friedman. Spatial comparison between wall shear stress measures and porcine arterial endothelial permeability. American Journal of Physiology-Heart and Circulatory Physiology, 286(5):H1916–H1922, 2004. doi: 10.1152/ajpheart.00897.2003.

CW Hirt, AA Amsden, and JL Cook. An arbitrary lagrangian–eulerian computing method for all flow speeds. Journal of computational physics, 135(2):203–216, 1997. doi: 10.1006/jcph.1997.5702.

Daniel Hirtler, Julio Garcia, Alex J Barker, and Julia Geiger. Assessment of intracardiac flow and vorticity in the right heart of patients after repair of tetralogy of fallot by flow-sensitive 4d mri. European radiology, 26: 3598–3607, 2016. doi: 10.1007/s00330-015-4186-1.

Ashifa Hudani, Safia Ihsan Ali, David Patton, Kimberley A Myers, Nowell M Fine, James A White, Steven Greenway, and Julio Garcia. 4d-flow mri characterization of pulmonary flow in repaired tetralogy of fallot. Applied Sciences, 13(5):2810, 2023. doi: 10.3390/app13052810.

Shelby Kutty, Quanliang Shang, Navya Joseph, Johannes T Kowallick, Andreas Schuster, Michael Steinmetz, David A Danford, Phillip Beerbaum, and Samir Sarikouch. Abnormal right atrial performance in repaired tetralogy of fallot: A cmr feature tracking analysis. International journal of cardiology, 248:136–142, 2017. doi: 10.1016/j.ijcard.2017.06.121.

Yue-Hin Loke, Francesco Capuano, Elias Balaras, and Laura J Olivieri. Computational modeling of right ventricular motion and intracardiac flow in repaired tetralogy of fallot. Cardiovascular engineering and technology, 13(1): 41–54, 2022. doi: 10.1007/s13239-021-00558-3.

Yue-Hin Loke, Ibrahim N Yildiran, Francesco Capuano, Elias Balaras, and Laura Olivieri. Tetralogy of fallot regurgitation energetics and kinetics: an intracardiac flow analysis of the right ventricle using computational fluid dynamics. The International Journal of Cardiovascular Imaging, pages 1–13, 2024. doi: 10.1007/s10554-024-03084-0.

PC Lu, HC Lai, and JS Liu. A reevaluation and discussion on the threshold limit for hemolysis in a turbulent shear flow. Journal of biomechanics, 34(10):1361–1364, 2001. doi: 10.1016/S0021-9290(01)00084-7.

JO Mangual, F Domenichini, and Gianni Pedrizzetti. Describing the highly three dimensional right ventricle flow. Annals of biomedical engineering, 40(8):1790–1801, 2012. doi: 10.1007/s10439-012-0540-5.

Guy Mareels, Dirk S De Wachter, and Pascal R Verdonck. Computational fluid dynamics-analysis of the niagara hemodialysis catheter in a right heart model. Artificial Organs, 28(7): 639–648, 2004. doi: 10.1111/j.1525-1594.2004.07371.x.

Amanda Mikhail, Giuseppe Di Labbio, Ahmed Darwish, and Lyes Kadem. How pulmonary valve regurgitation after tetralogy of fallot repair changes the flow dynamics in the right ventricle: An in vitro study. Medical Engineering & Physics, 83:48–55, 2020. doi: 10.1016/j.medengphy.2020.07.014.

Christopher Naoum, Philipp Blanke, João L Cavalcante, and Jonathon Leipsic. Cardiac computed tomography and magnetic resonance imaging in the evaluation of mitral and tricuspid valve disease: implications for transcatheter interventions. Circulation: Cardiovascular Imaging, 10(3):e005331, 2017. doi: 10.1161/CIRCIMAGING.116.005331.

Franck Nicoud, Hubert Baya Toda, Olivier Cabrit, Sanjeeb Bose, and Jungil Lee. Using singular values to build a subgrid-scale model for large eddy simulations. Physics of fluids, 23(8): 085106, 2011. doi: 10.1063/1.3623274.

Fabio Nobile and Luca Formaggia. A stability analysis for the arbitrary lagrangian eulerian formulation with finite elements. East-West Journal of Numerical Mathematics, 7:105–132, 1999.

Achilles J Pappano and Withrow Gil Wier. Cardiovascular physiology-e-book. Elsevier Health Sciences, 2018. ISBN 978-0-323-08697-4.

Louis P Parker, Anders Svensson Marcial, Torkel B Brismar, Lars Mikael Broman, and Lisa Prahl Wittberg. Impact of altered vena cava flow rates on right atrium flow characteristics. Journal of Applied Physiology, 132(5): 1167–1178, 2022. doi: 10.1152/japplphysiol.00649.2021.

Stephen B Pope. Turbulent flows, volume 12. 2001. doi: 10.1088/0957-0233/12/11/705.

Alfio Quarteroni, Andrea Manzoni, Christian Vergara, et al. Mathematical modelling of the human cardiovascular system: data, numerical approximation, clinical applications, volume 33. Cambridge University Press, 2019. ISBN 9781108480390.

Andrew N Redington. Determinants and assessment of pulmonary regurgitation in tetralogy of fallot: practice and pitfalls. Cardiology clinics, 24(4):631–639, 2006. doi: 10.1016/j.ccl.2006.08.007.

Francesca Renzi, Christian Vergara, Marco Fedele, Vincenzo Giambruno, Alfio Maria Quarteroni, Giovanni Puppini, and Giovanni Battista Luciani. Accurate and efficient 3d reconstruction of right heart shape and motion from multi-series cine-mri. bioRxiv, pages 2023–06, 2023. doi: 10.1101/2023.06.28.546872.

Gerald J Riccardello Jr, Darshan N Shastri, Abhinav R Changa, Kiran G Thomas, Max Roman, Charles J Prestigiacomo, and Chirag D Gandhi. Influence of relative residence time on side-wall aneurysm inception. Neurosurgery, 83(3):574–581, 2018. doi: 10.1093/neuros/nyx433.

Federica Sacco, Bruno Paun, Oriol Lehmkuhl, Tinen L Iles, Paul A Iaizzo, Guillaume Houzeaux, Mariano Vázquez, Constantine Butakoff, and Jazmin Aguado-Sierra. Evaluating the roles of detailed endocardial structures on right ventricular haemodynamics by means of cfd simulations. International journal for numerical methods in biomedical engineering, 34(9):e3115, 2018. doi: 10.1002/cnm.3115.

Selvi Senthilnathan, Andreea Dragulescu, and Luc Mertens. Pulmonary regurgitation after tetralogy of fallot repair: a diagnostic and therapeutic challenge. Journal of Cardiovascular Echography, 23(1):1–9, 2013. doi: 10.4103/2211-4122.117975.

Jung Hee Seo, Vijay Vedula, Theodore Abraham, Albert C Lardo, Fady Dawoud, Hongchang Luo, and Rajat Mittal. Effect of the mitral valve on diastolic flow patterns. Physics of fluids, 26(12), 2014. doi: 10.1063/1.4904094.

Keith Stein, Tayfun Tezduyar, and Richard Benney. Mesh moving techniques for fluid-structure interactions with large displacements. J. Appl. Mech., 70(1):58–63, 2003. doi: 10.1115/1.1530635.

Simone Stella, Christian Vergara, Luca Giovannacci, Alfio Quarteroni, and Giorgio Prouse. Assessing the disturbed flow and the transition to turbulence in the arteriovenous fistula. Journal of biomechanical engineering, 141(10): 101010, 2019. doi: 10.1115/1.4043448.

Marco Stevanella, Francesco Maffessanti, Carlo A Conti, Emiliano Votta, Alice Arnoldi, Massimo Lombardi, Oberdan Parodi, Enrico G Caiani, and Alberto Redaelli. Mitral valve patient-specific finite element modeling from cardiac mri: application to an annuloplasty procedure. Cardiovascular Engineering and Technology, 2: 66–76, 2011. doi: 10.1007/s13239-010-0032-4.

Shin-ichiro Sugiyama, Kuniyasu Niizuma, Toshio Nakayama, Hiroaki Shimizu, Hidenori Endo, Takashi Inoue, Miki Fujimura, Makoto Ohta, Akira Takahashi, and Teiji Tominaga. Relative residence time prolongation in intracranial aneurysms: a possible association with atherosclerosis. Neurosurgery, 73(5):767–776, 2013. doi: 10.1227/NEU.0000000000000096.

Dalin Tang, Chun Yang, Tal Geva, and J Pedro. Image-based patient-specific ventricle models with fluid–structure interaction for cardiac function assessment and surgical design optimization. Progress in pediatric cardiology, 30 (1-2):51–62, 2010. doi: 10.1016/j.ppedcard.2010.09.007.

Tayfun E Tezduyar and Masayoshi Senga. Stabilization and shock-capturing parameters in supg formulation of compressible flows. Computer methods in applied mechanics and engineering, 195(13-16):1621–1632, 2006. doi: 10.1016/j.cma.2005.05.032.

Alexandre This, Ludovic Boilevin-Kayl, Miguel A Fernández, and Jean-Frédéric Gerbeau. Augmented resistive immersed surfaces valve model for the simulation of cardiac hemodynamics with isovolumetric phases. International journal for numerical methods in biomedical engineering, 36(3):e3223, 2020. doi: 10.1002/cnm.3223.

Emanuela R Valsangiacomo Buechel and Luc L Mertens. Imaging the right heart: the use of integrated multimodality imaging. European heart journal, 33(8):949–960, 2012. doi: 10.1093/eurheartj/ehr490.

Christian Vergara, Davide Le Van, Maurizio Quadrio, Luca Formaggia, and Maurizio Domanin. Large eddy simuations of blood dynamics in abdominal aortic aneurysms. Medical Engineering & Physics, 47:38–46, 2017. doi: 10.1016/j.medengphy.2017.06.030.

Francesco Viola, Vamsi Spandan, Valentina Meschini, Joshua Romero, Massimiliano Fatica, Marco D de Tullio, and Roberto Verzicco. Fseigpu: Gpu accelerated simulations of the fluid– structure–electrophysiology interaction in the left heart. Computer physics communications, 273:108248, 2022. doi: 10.1016/j.cpc.2021.108248.

Emiliano Votta, Enrico Caiani, Federico Veronesi, Monica Soncini, Franco Maria Montevecchi, and Alberto Redaelli. Mitral valve finite-element modelling from ultrasound data: a pilot study for a new approach to understand mitral function and clinical scenarios. Philosophical Transactions of the Royal Society A: Mathematical, Physical and Engineering Sciences, 366(1879): 3411–3434, 2008. doi: 10.1098/rsta.2008.0095.

Hadi Wiputra, Chang Quan Lai, Guat Ling Lim, Joel Jia Wei Heng, Lan Guo, Sanah Merchant Soomar, Hwa Liang Leo, Arijit Biwas, Citra Nurfarah Zaini Mattar, and Choon Hwai Yap. Fluid mechanics of human fetal right ventricles from image-based computational fluid dynamics using 4d clinical ultrasound scans. American Journal of Physiology-Heart and Circulatory Physiology, 311(6):H1498–H1508, 2016. doi: 10.1152/ajpheart.00400.2016.

Hadi Wiputra, Ching Kit Chen, Elias Talbi, Guat Ling Lim, Sanah Merchant Soomar, Arijit Biswas, Citra Nurfarah Zaini Mattar, David Bark, Hwa Liang Leo, and Choon Hwai Yap. Human fetal hearts with tetralogy of fallot have altered fluid dynamics and forces. American Journal of Physiology-Heart and Circulatory Physiology, 315(6):H1649–H1659, 2018. doi: 10.1152/ajpheart.00235.2018.

Ajit P Yoganathan, Edward G Cape, Hsing-Wen Sung, Frank P Williams, and Abdul Jimoh. Review of hydrodynamic principles for the cardiologist: applications to the study of blood flow and jets by imaging techniques. Journal of the American College of Cardiology, 12(5): 1344–1353, 1988. doi: 10.1016/0735-1097(88)92620-4.

Alberto Zingaro, Michele Bucelli, Roberto Piersanti, Francesco Regazzoni, Alfio Quarteroni, et al. An electromechanics-driven fluid dynamics model for the simulation of the whole human heart. Journal of Computational Physics, 504: 112885, 2024. doi: 10.1016/j.jcp.2024.112885.

